# Post-injury born oligodendrocytes integrate into the glial scar and inhibit growth of regenerating axons by premature myelination

**DOI:** 10.1101/2021.10.15.464557

**Authors:** Jian Xing, Bruce A. Rheaume, Juhwan Kim, Agnieszka Lukomska, Muhammad S. Sajid, Ashiti Damania, Ephraim F. Trakhtenberg

**Affiliations:** Department of Neuroscience, University of Connecticut School of Medicine, 263 Farmington Ave., Farmington, CT, 06030, USA

## Abstract

Pathologies of the central nervous system (CNS) white matter often result in permanent functional deficits because mature mammalian projection neurons fail to regenerate long-distance axons after injury. A major barrier to axonal regenerative research is that the CNS axons that regenerate in response to experimental treatments stall growth before reaching their post-synaptic targets. Here, we test the hypothesis that premature, *de novo*, myelination of regenerating axons stalls their growth, even after bypassing the glial scar. To test this hypothesis, first, we used single cell RNA-seq (scRNA-seq) and immunohistological analysis to investigate whether post-injury born oligodendrocytes integrate into the glial scar after optic nerve injury. Then, we used a multiple sclerosis model of demyelination concurrently with the stimulation of axon regeneration by Pten knockdown (KD) in projection neurons after optic nerve injury. We found that post-injury born oligodendrocytes integrate into the glial scar, where they are susceptible to the demyelination treatment, which prevented premature myelination, and thereby enhanced Pten KD-stimulated axon regeneration. We also present a website for comparing the gene expression of scRNA-seq-profiled optic nerve oligodendrocytes under physiological and pathophysiological conditions.

**SIGNIFICANCE STATEMENT:** Myelin debris from degenerating axons along with reactive astrocytes in the glial scar inhibit CNS axon regeneration. However, even with the recently developed experimental approaches which activate axons to regenerate passed the glial scar, almost all axons still stall growth before reaching their post-synaptic targets. Here, we show that post-injury born oligodendrocytes integrate into the glial scar, and that other than myelin debris, live oligodendrocytes prematurely myelinating the regenerating axons inhibit growth, even if the axons have already regenerated passed the glial scar.

## INTRODUCTION

Mammalian central nervous system (CNS) projection neurons’ failure to regenerate axons disrupted by white matter injury to the brain, spinal cord, or optic nerve results in permanent disabilities. Like other CNS projection neurons, retinal ganglion cells (RGCs) do not regenerate axons disrupted by optic nerve crush (ONC)^1,2^. Because molecules found to regulate regeneration of RGC axons, such as Pten and Klf7^3,4^, also affect spinal cord regeneration^4,5^, the mechanisms of axonal regeneration may be similar across CNS projection neurons, while the mechanisms of their pathway-finding vary. A number of intracellular and extracellular factors have been discovered to affect axon regeneration (as reviewed elsewhere, e.g.,^6–12^, but even with manipulation of potent tumorigenic factors <1% of axons regenerated the full-length to their post-synaptic targets^13–15^. Thus, although stimulating neuronal intrinsic mechanisms of axon regeneration was sufficient for bypassing extracellular inhibitors of axon growth that are associated with the glial scar and myelin (e.g., CSPG, MAG, NogoA, OMgp, Semaphorins)^10,16^, almost all regenerating axons stall growth far before reaching targets in the brain.

Here, we investigated why the regeneration of axons that respond to regenerative treatments stalls (e.g.,^13–15,17–22^). We hypothesized that premature *de novo* myelination of the axons that are experimentally-stimulated to regenerate stalls their growth even after the axons bypass the glial scar and grow over pre-injury myelin. During CNS development, axon myelination is delayed until the axons have reached their postsynaptic targets in the brain. For example, myelination of the optic nerve axons in mice begins on postnatal day 7 – over a week after these axons have reached their targets in the brain^23,24^. Then, myelination prevents sprouting and stabilizes axons during the developmental pruning period^25^. However, optic nerve axons experimentally-induced to regenerate after injury are myelinated even while they are still growing (and eventually the growth stalls)^26^. Thus, it is possible that primarily the premature *de novo* myelination, rather than the pre-existing myelin and its debris, is what stalls experimental regeneration even after axons have bypassed the inhibitory glial scar. Indeed, conditional deletion in oligodendrocytes of the neurological diseases-associated brain-specific angiogenesis inhibitors (BAIs; which bind to the axon growth-inhibitory NogoA reticulon-4 receptor; Rtn4R), was recently shown to rescue axonal growth of co-cultured neurons^27^, further supporting our hypothesis that live oligodendrocytes, and not only their debris after injury, could inhibit axon regeneration. To test our hypothesis, we used single cell RNA-seq (scRNA-seq) to analyze the effects of ONC on optic nerve oligodendrocytes, and a cuprizone-induced chronic demyelination (that injures oligodendrocytes to model multiple sclerosis) during traumatic optic neuropathy (modeled by ONC), which acutely and irreversibly disrupts the neuronal axons while modifying the extra-axonal tissue environment^2^.

## RESULTS

### Combining models of multiple sclerosis demyelination and traumatic optic nerve injury

In the multiple sclerosis model, a cuprizone diet injures or kills mature oligodendrocytes^28,29^. Because the optic nerve is amongst the least susceptible CNS regions to cuprizone-induced demyelination^30,31^, we added rapamycin injections, which reduce spontaneous remyelination, resulting in an overall greater level of demyelination^32,33^. After the demyelination regimen is stopped, both recovered mature and newly-formed oligodendrocytes remyelinate axons^32,33^. Because rapamycin may reduce axon regeneration after injury^34^, we discontinued rapamycin injections two weeks prior to ONC^35,36^ (experimental timeline in **Fig. 1A**). Because remyelination only becomes detectable about a week after discontinuation of the cuprizone diet^32,33^, we stopped the diet one week prior to euthanizing the animals. After ONC, RGC-specific knockdown (KD) of Pten (see Methods and **Supplementary Figure 1**) was used to stimulate a subset of RGCs to re-grow axons past the glial scar and over myelin^2,18,34^.

**Fig. 1.**
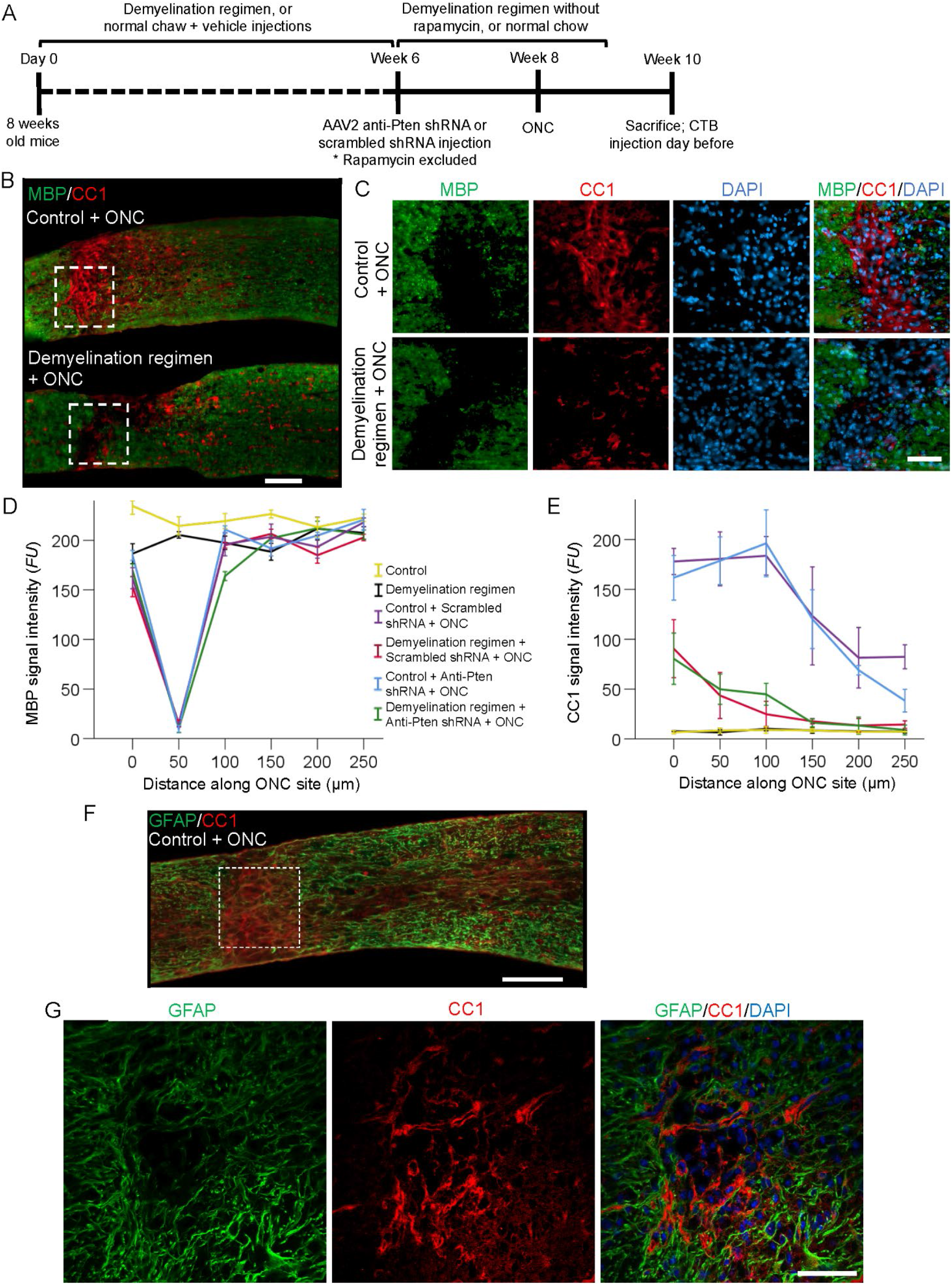
Loss of myelin in the ONC site and the effects of demyelination regimen on the optic nerve. (**A**) Experimental timeline. At 6 weeks after starting the demyelination regimen, AAV2 expressing anti-Pten or scrambled control shRNA was injected intravitreally, 2 weeks later ONC was performed, and mice were sacrificed at either 2, 4, or 6 weeks after that. CTB (axonal tracer) was injected intravitreally 1 day before sacrifice. Rapamycin was excluded from the demyelination regimen 6 weeks after starting the regimen, and the remaining regiment stopped 1 week prior to sacrifice. For the experiments shown in Figs. 3 and 7, mice were sacrificed 6 or 2 weeks after injury, respectively, and viruses and CTB were injected as above, without accompanying demyelination regimen, but in Fig. 7 also with intravitreal cuprizone or saline injection right after ONC in 10-week old mice. Mice sacrificed at 4 weeks after ONC were used for Fig. 6. (**B-C**) Representative images of injured optic nerves longitudinal sections immunostained for CC1 (mature oligodendrocyte marker), MBP (myelin marker), and DAPI (nuclear marker), either without (upper panels) or with (lower panels) demyelination regimen. Injury site outlined with dashed white lines box. Scale bars: 100 μm main panels (*B*), 50 μm insets (*C*). (**D-E**) Quantification of MBP (*D*) and CC1 (*E*) immunofluorescence signal intensity, represented in fluorescent units (*FU*s), at increasing distances along the optic nerve injury site (the region devoid of MBP and its surroundings, as shown in panels *B-C*) or along an equivalent uninjured region, across conditions as marked. Representative images for all the conditions shown in Supplementary Figure 3 (MBP) and Supplementary Figure 6 (CC1). Mean ± S.E.M shown; *n* = 4 cases for each condition. ANOVA with repeated measures, sphericity assumed, overall *F* = 35.0 (MBP) and *F* = 3.5 (CC1), *p* < 0.001 (MBP) and *p* < 0.001 (CC1); *p*-values of pairwise comparisons by posthoc LSD are shown in Table 1A (MBP) and 1B (CC1). (**F**) Representative images of the longitudinal optic nerve sections immunostained for glial fibrillary acidic protein (GFAP; labels astrocytes) and CC1 2 weeks after ONC. Injury site outlined with dashed white lines box (shown in *F*). Scale bar, 100 μm. (**G**) Inset: Confocal images of the injury site show cellular segregation between GFAP and CC1 signals; counterstained with DAPI to label nuclei. Scale bar, 50 μm.

### Demyelination regimen remodels the glial scar after traumatic optic nerve injury

The demyelination regimen led to robust loss of myelin in the corpus callosum, as shown by a decrease of myelin basic protein (MBP)^37^ along the corpus callosum tracts (**Supplementary Figure 2A,C**). Demyelination in the corpus callosum was accompanied by a reduction in oligodendrocytes, whose soma is labeled by immunostaining for the CC1 marker of mature oligodendrocytes^38–40^ (**Supplementary Figure 2B,D**). A greater loss of myelin in the corpus callosum compared to the optic nerve after demyelination diet was previously attributed to the optic nerve being more resistant to cuprizone^30,31,41^. We also observed only a 10% (*p* < 0.01) decrease in the MBP signal after the cuprizone diet in the uninjured optic nerve, as well as in the uninjured segment of the injured optic nerve distal from the ONC site (where the disconnected by the injury axonal segments are degenerating). However, even without a demyelination regimen, loss of MBP signal was apparent within the ONC site (where the glial scar forms; **Supplementary Figure 3, Fig. 1B-D**, and **Table 1A**). It is possible that the differences in efficiency of myelin debris clearance between corpus callosum and the optic nerve may underlie their differential response to the cuprizone diet.

**Table 1.** Pairwise comparison *p*-values for Fig. 1D and Supplementary Figure 3 (MBP), as well as for Fig. 1E, and Supplementary Figure 6 (CC1). Pairwise comparisons between the conditions for MBP signal intensity (*A*) and CC1 signal intensity (*B*) were performed by ANOVA with repeated measures and posthoc LSD. The *p*-values for each nonredundant comparison are shown, and significant differences (*p* ≤ 0.05) indicated by an asterisk (*).

| **A** | Demyelin-<br>ation regi-<br>men | Control +<br>Scrambled<br>shRNA + ONC | Demyelination<br>regimen +<br>Scrambled<br>shRNA + ONC | Control +<br>Anti-Pten<br>shRNA +<br>ONC | Demyelination<br>regimen + Anti-Pten shRNA +<br>ONC |
| --- | --- | --- | --- | --- | --- |
| Control | 0.00 * | 0.02 * | 0.01 * | 0.00 * | 0.00 * |
| Demyelination regimen |  | 0.06 | 0.03 * | 0.02 * | 0.02 * |
| Control + Scrambled<br>shRNA + ONC |  |  | 0.12 | 0.35 | 0.43 |
| Demyelination regimen +<br>Scrambled shRNA + ONC |  |  |  | 0.07 | 0.69 |
| Control + Anti-Pten<br>shRNA + ONC |  |  |  |  | 0.01 * |

**Table 1.** Pairwise comparison *p*-values for Fig.
| **B** | Demyelin-<br>ation regi-<br>men | Control +<br>Scrambled<br>shRNA + ONC | Demyelination<br>regimen +<br>Scrambled<br>shRNA + ONC | Control +<br>Anti-Pten<br>shRNA +<br>ONC | Demyelination<br>regimen + Anti-Pten shRNA +<br>ONC |
| --- | --- | --- | --- | --- | --- |
| Control | 0.96 | 0.02 * | 0.11 | 0.02 * | 0.14 |
| Demyelination regimen |  | 0.01 * | 0.12 | 0.02 * | 0.12 |
| Control + Scrambled<br>shRNA + ONC |  |  | 0.01 * | 0.64 | 0.00 * |
| Demyelination regimen +<br>Scrambled shRNA + ONC |  |  |  | 0.05 * | 0.89 |
| Control + Anti-Pten<br>shRNA + ONC |  |  |  |  | 0.02 * |

Initially, by 3 days after ONC, the injury causes decrease in CC1 signal in the injury site (due to the death of oligodendrocytes), and an increase in the MBP signal (due to the accumulation of myelin debris from the dead oligodendrocytes) (**Supplementary Figure 4**). Apparent loss of myelin in the optic nerve injury site by 2 weeks after ONC (**Fig. 1B-D**), however, is due to the phagocytosing immune cells concentrating there to clear debris^8,42^. These immune cells do not appear to invade more distal nerve regions where axons undergo slow Wallerian degeneration^43^. Accordingly, using scRNA-seq analysis, we found a 14-fold increase in the immune cell population (Iba1^44^, CD45^45^, and C1qc^46^ positive cells cluster) in the injury site 2 weeks after ONC (**Fig. 2A-E** and **Supplementary Figure 5A-C**), consistent with an increase in the density of cells with normal DAPI morphology in the injury site by 2 weeks after ONC (**Fig. 1C** and **Supplementary Figure 3**), as the immune cells infiltrate the injury site^8^ and clear myelin debris^42^. Spontaneous clearance of myelin debris from within the injury site suggests that they are not major constituents of the persisting optic nerve glial scar, however, the remaining myelin sheath still wrapping the degenerating axons by the injury site, along with myelin debris at the boundary of the glial scar, may hinder axon regeneration^10,16^. Although the effect of the demyelination regimen on pre-existing myelin and its debris in the optic nerve is modest, we hypothesized that the regimen would be sufficient to block *de novo* myelination of the regenerating axons, as it is well-established to injure oligodendrocytes^28,29^ even if their pre-existing myelin is only partially cleared, and it thus may reduce and delay the formation of new myelin, thereby permitting more axon regeneration.

**Fig 2.**
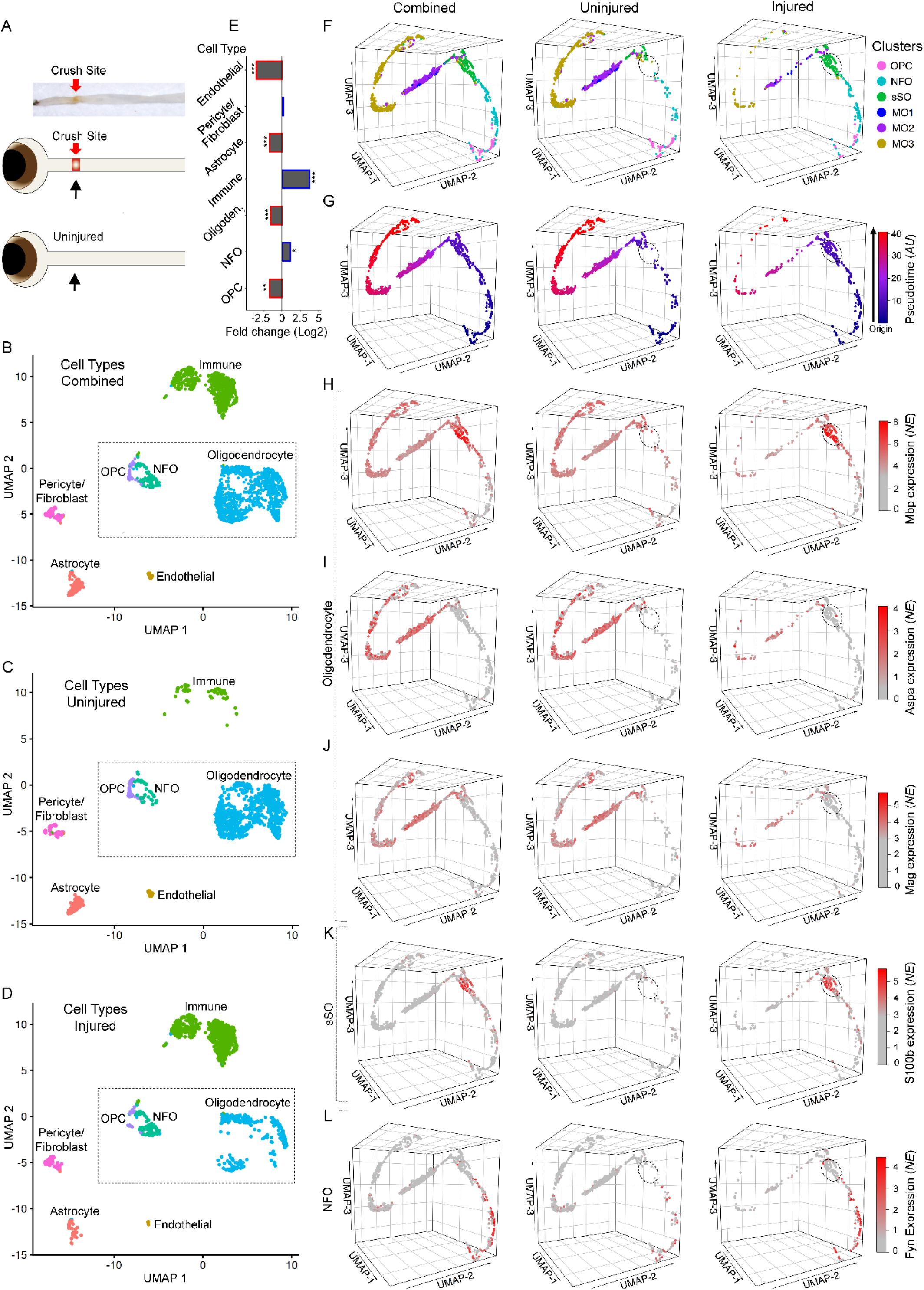
Single cell transcriptome profiling of oligodendrocyte lineage in the uninjured and injured optic nerves. (**A**) Experimental design: (*Top panel*) a representative image of a resected optic nerve with red arrow indicating the crush site from where the tissue samples were dissected for scRNA-seq; (*bottom panels*) a cartoon representation of the injured and uninjured samples analyzed by scRNA-seq. (**B-D**) 2-dimensional Uniform Manifold Approximation and Projection (UMAP) of the scRNA-seq-identified cell types from the injured and uninjured optic nerve tissues, combined (in *B*), uninjured only (in *C*), and injured only (in *D*). The groups of cells comprising the specific cell types are color-coded and annotated as marked. The UMAPs of the respective markers’ expression used to determine the cell types are shown in the Supplementary Figure 5. The dashed box indicates oligodendrocyte lineage cells for downstream analyses (in *F*-*L*). (**E**) Fold changes after ONC injury in the populations of cell types comprising the optic nerve tissue, as marked. Fold change significance determined by EdgeR analysis, as indicated (\*\*\**p* < 0.001, \*\**p* < 0.01, and \**p* < 0.05; see Methods). (**F**) Cluster analysis (by the Seurat algorithm, see Methods) of the cell types comprising oligodendrocyte lineage (OPC, NFO, and Oligodendrocytes; outlined by dashed box in *B*-*D*), represented by 3-dimensional (3D) UMAPs for better visualization of all cells combined, or uninjured only, or injured only. Clusters comprising specific cell types are color-coded and annotated, as marked (OPC = oligodendrocyte progenitor cell, NFO = newly-formed oligodendrocyte, sSO = small spidery oligodendrocyte, MO = mature oligodendrocyte; cluster sSO is encircled in the UMAPs of uninjured and injured cells in *E*-*L* for better visualization). (**G**) 3D UMAPs of cells comprising oligodendrocyte lineage (as in *F*; with all cells combined, or uninjured only, or injured only), color-coded based on psudotimeline analysis (by the Monocle3 algorithm; *AU* = Arbitrary units; see Methods), which represents the relative transcriptomic changes progressing along the trajectory of OPCs differentiating into NFOs, then developing into sSOs, and finally maturing into MOs. (**H-L**) 3D UMAPs of cells comprising oligodendrocyte lineage (as in *F-G*; with all cells combined, or uninjured only, or injured only), showing the expression of respective cell types’ gene markers (as marked): MBP (MO and sSO, in *H*), Aspa (MO, in *I*), MAG (MO, in *J*), S100β (sSO, in *K*), Fyn (NFO, in *L*), and Pdgfra (OPC; due to space limit here shown in the Supplementary Figure 5M). Scale bar, color-coded normalized expression (*NE*).

We then further characterized the injury site and expectedly found astrocytes (labeled with the glial fibrillary acidic protein, GFAP) surrounding the injury site (**Fig. 1F-G**). Within the injury site, we found CC1+/MBP− oligodendrocytes with the morphology of newly-formed oligodendrocytes (NFOs), including maturing small spidery oligodendrocytes (sSO)^47^ that are primed-to-myelinate axons (if axons would be available in proximity)^48,49^ (**Fig. 1, Supplementary Figures 3, 6**, and **Table 1**). Within the ONC site, CC1+ oligodendrocytes were negative for another oligodendrocyte marker, Aspa^50^, but were positive for Aspa in the uninjured region of the optic nerve (**Supplementary Figure 7**). Oligodendrocyte progenitor cells (OPC) are not labeled by the CC1 marker but are present by the glial scar after injury^51^ and differentiate into CC1+ oligodendrocytes^52^. The presence of the post-injury-born CC1+/MBP−/Aspa− oligodendrocytes within the glial scar, however, was reduced by the demyelination regimen, whereas the density of oligodendrocytes did not appear to change in the non-crushed segments of the optic nerve (**Fig. 1B-E, Supplementary Figure 6**, and **Table 1B**).

### Changes in oligodendrocyte subpopulations in the glial scar revealed by scRNA-seq

These data are consistent with the scRNA-seq analysis of the injury site after ONC, which showed a 3-fold decrease in the already small (relative to other optic nerve cell types) population of Pdgfra+ OPCs^53^, along with a 3-fold decrease in the mature oligodendrocyte population (**Fig. 2A-E** and **Supplementary Figure 5D-G**). However, in the injury site after ONC, Fyn+ NFOs^54,55^ increased 2-fold (**Fig. 2A-E** and **Supplementary Figure 5H**), and MBP+/S100β+/MAG−/Aspa− oligodendrocytes primed to myelinate axons emerged (encircled in **Fig. 2** panels **F-K**). These post-injury-emerging oligodendrocytes are negative for the markers of fully mature oligodendrocytes, such as MAG and Aspa, but are positive for S100β, which (in addition to astrocytes) is expressed in OPCs and maturing NFOs until they contact axons and consequently downregulate S100β as they progress towards full maturation^56,57^ (**Fig. 2F-K**). These post-injury-emerging oligodendrocytes express MBP mRNA (encircled in **Fig. 2H**), consistent with the CC1+ maturing sSOs accumulating MBP mRNA prior to beginning its translation into protein, which becomes detectible by immunostaining at a later stage of oligodendrocytes maturation^47^, and thus they were also negative for MBP protein by immunostaining in the injury site (**Fig. 1B-E, Supplementary Figures 3, 6**, and **Table 1**). The post-injury-emerging sSOs bioinformatically grouped into a discrete subpopulation of developing oligodendrocytes (green cluster labeled sSO in **Fig. 2F**), that was absent in the uninjured optic nerve. A small fraction of the fully mature MBP+/S100β−/MAG+/Aspa+ oligodendrocytes is also present in the injury site (**Fig. 2F-K**). Accordingly, the psudotimeline trajectory shows the progression from OPCs into NFOs and then into the primed-to-myelinate MBP_mRNA_+:_protein_−/S100β+/MAG−/Aspa− sSOs, and finally into the fully mature MBP+/S100β−/MAG+/Aspa+ oligodendrocytes (**Fig. 2F-G**). In the uninjured optic nerve, there are 2-fold less NFOs, and the intermediate MBP_mRNA_+:_protein_−/S100β+/MAG−/Aspa− sSO subpopulation of oligodendrocytes is nearly absent compared to the injured optic nerve. These data suggest that the different subtypes of numerous fully mature oligodendrocytes (MOs) in the uninjured optic nerve are not recently-born, consistent with the low physiological turnover rate of the oligodendrocytes in the optic nerve^58,59^ and the segregation of MOs into subtypes in other CNS regions^60^. In the injury site, however, this small subpopulation of the fully mature MBP+/S100β−/MAG+/Aspa+ oligodendrocytes include mostly post-injury-born that matured fully and only some of those that survived the injury (**Fig. 2F-K**). This is expected because shortly after injury (3 days after ONC) fewer surviving oligodendrocytes are detected (**Supplementary Figure 4**), whereas at 2 weeks after injury more oligodendrocytes are detected, compared to uninjured optic nerve, indicating that most oligodendrocytes in the injury site by 2 weeks after ONC are newly-born (**Fig. 1B-D**). Taken together, the immunohistological and scRNA-seq data suggest a decrease in the MO population, with only a small fraction of the injured oligodendrocytes surviving, an increase in the NFOs, and the emergence of post-injury born primed-to-myelinate oligodendrocytes by 2 weeks after ONC. Thus, post-injury-born and the surviving-injured oligodendrocytes contribute to the glial scar, where they are more susceptible to the demyelination regimen, compared to those distal to the injury site located in the uninjured environment.

### Gene Browser of oligodendrocyte subtypes under physiological and pathophysiological conditions

We also developed the Subtypes Gene Browser web application (in the same format we previously presented for retinal ganglion cell subtypes^61^), which provides a platform for analyzing and comparing gene expression profiles in OPC and oligodendrocyte subtypes from the uninjured and injured optic nerves (https://health.uconn.edu/neuroregeneration-lab/subtypes-gene-browser). The browser offers three types of analyses for any gene in the OPC and oligodendrocyte subtypes: Means and SEM^62^, Violin Plot of cell density for different gene expression levels^63^, and Box Plot for comparing medians using the Tukey Box-and-Whisker plot with whiskers set to 1.5 times the interquartile range (IQR)^64–66^.

### Demyelination regimen enhances experimental axon regeneration

Pre-injury myelin is cleared from the injury site, even in untreated animals, to the extent that MBP is barely detectable by immunostaining, while oligodendrocytes are present in the injury site (by 2 weeks after injury; **Fig. 1B-D, Supplementary Figure 3**, and **Table 1A**). Therefore, such an environment as in the ONC injury site provides a unique opportunity to visualize, by immunostaining, whether the axons experimentally-stimulated to regenerate are *de novo* myelinated before reaching their post-synaptic targets in the brain. We previously showed that axons, experimentally-stimulated to regenerate, stall growth before reaching their targets, even at 6 weeks after injury^15^. Thus, to allow new myelin to accumulate sufficiently for detection by immunostaining for MBP, we analyzed the injury site 6 weeks after ONC and Pten KD-stimulated axon regeneration. Consistent with prior reports^26^, confocal analysis showed the MBP signal associated with regenerating axons in the injury site, where MBP detection by immunostaining is otherwise bordering noise (**Fig. 3**).

**Fig. 3.**
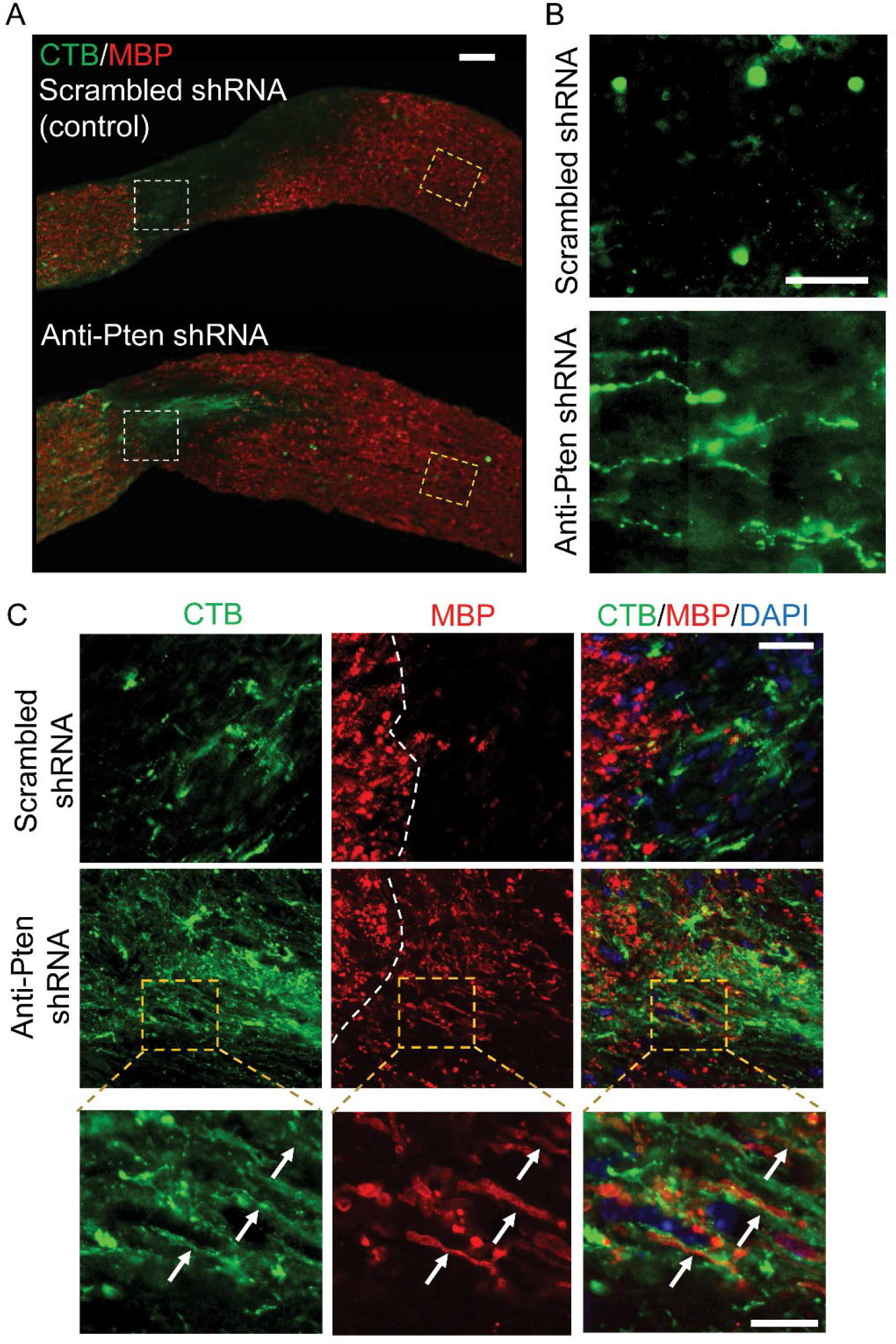
Regenerating axons become myelinated prematurely. (**A**) Representative images of the optic nerve longitudinal sections immunostained for CTB and MBP at 6 weeks after ONC, either pre-treated with AAV2 expressing scrambled shRNA (control) or anti-Pten shRNA. Injury site outlined with dashed white lines box. Scale bar, 100 μm. (**B**) Zoomed-in insets (outlined in yellow dashed lines in *A*) show termination of stalled Pten KD-stimulated regenerating axons at longer distances (1.5 mm from the ONC site; *lower panel*), which are otherwise masked by merged MBP signal in panel *A*. No regenerating axons in control (*upper panel*). Exposure in insets digitally increased for better visualization of the regenerating axons. Scale bar, 50 μm. (**C**) Confocal images of the insets (outlined in white dashed lines in *A*) show myelination of Pten KD-stimulated regenerating axons growing through the injury site (*lower panels*). No regenerating axons or MBP in control injury site (*upper panels*). Dashed white line demarcates beginning of the injury site (to the right). Insets below show co-localization of MBP with regenerating axons visualized by CTB (*arrows*). Scale bars, 50 μm (main panels), 20 μm (insets).

Next, we evaluated the effect of the demyelination regimen on axon regeneration and found significantly longer Pten KD-stimulated regenerating axons compared to Pten KD alone, and even without Pten KD, some axons regenerated, albeit over a short distance (**Fig. 4A-B** and **Table 2A**). There was also a modest positive effect on RGC survival^15,67^ in the Pten KD/cuprizone combination compared to Pten KD alone or control (**Fig. 5A,C** and **Table 3A**). In this assay, axon regeneration is typically assessed at 2 weeks after ONC, and RGCs that respond to Pten KD regenerate axons during this time window^15,18^ (experimental timeline in **Fig. 1A**). However, this time window was insufficient for allowing *de novo* myelination of nascent regenerated segments of the axons to a level that is unambiguously detectable by immunostaining for MBP. By immunostaining optic nerve sections, we found MBP is detectable on the regenerating axons without a cuprizone diet at 4 weeks after ONC (**Fig. 6A,C**). Although the MBP signal at 4 weeks was not as robust as at 6 weeks after ONC (**Fig. 3**), mice cannot be sustained on a cuprizone diet for that long because of its toxicity. Thus, as mice needed to undergo pre-treatment with the cuprizone diet in order to assess its effect on *de novo* myelination of the regenerating axons, we were only able to sustain mice on a cuprizone diet for up to 4 weeks after ONC. Using this approach, we found that the demyelination regimen indeed decreased *de novo* myelination of the regenerated axons (**Fig. 6**).

**Table 2.**
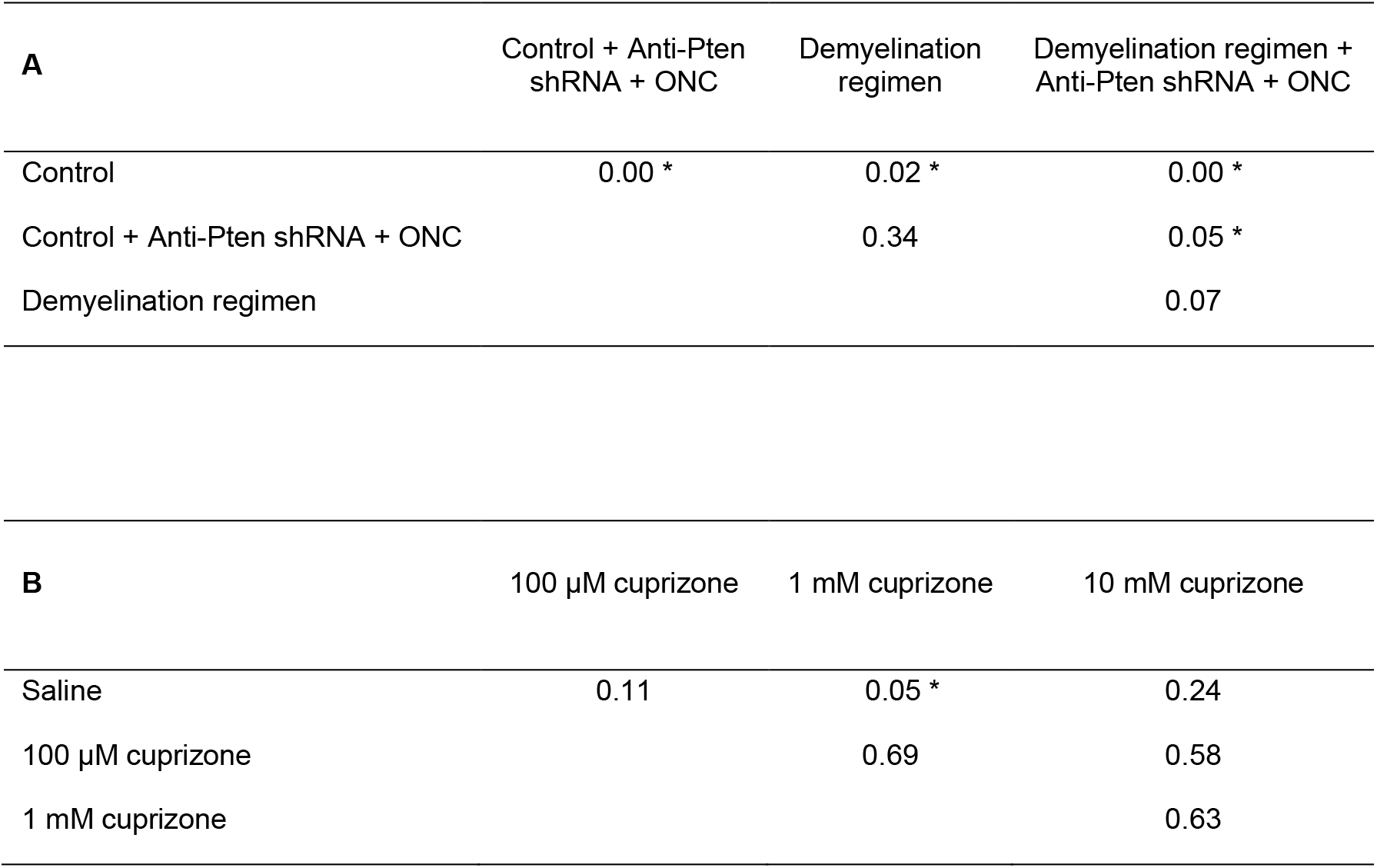
Pairwise comparison *p*-values for Figs. 4 and 7 on axon regeneration. Pairwise comparisons between the conditions for pre-treatment with the demyelination regimen (*A*) and post-injury treatment with intravitreal injections (*B*) were performed by ANOVA with repeated measures and posthoc LSD. The *p*-values for each nonredundant comparison are shown, and significant differences (*p* ≤ 0.05) indicated by an asterisk (*).

**Table 3.** Pairwise comparison *p*-values for Figs. 5C-D on RGC survival. Pairwise comparisons between the conditions for pre-treatment with the demyelination regimen (*A*) and post-injury treatment with intravitreal injections (*B*) were performed by ANOVA with repeated measures and posthoc LSD. The *p*-values for each nonredundant comparison are shown, and significant differences (*p* ≤ 0.05) indicated by an asterisk (*).

| **A** | Demyelination regimen +<br>Scrambled shRNA + ONC | Control + Anti-Pten<br>shRNA + ONC | Demyelination regimen +<br>Anti-Pten shRNA + ONC |
| --- | --- | --- | --- |
| Control + Scrambled<br>shRNA + ONC | 0.10 | 0.38 | 0.01 * |
| Demyelination regimen +<br>Scrambled shRNA + ONC |  | 0.37 | 0.09 |
| Control + Anti-Pten<br>shRNA + ONC |  |  | 0.02 * |

**Table 3.** Pairwise comparison *p*-values for Figs.
| **B** | 100 μM cuprizone | 1 mM cuprizone | 10 mM cuprizone |
| --- | --- | --- | --- |
| Saline | 0.21 | 0.24 | 0.21 |
| 100 μM cuprizone | | 0.93 | 1.00 |
| 1 mM cuprizone |  |  | 0.93 |

**Fig. 4.**
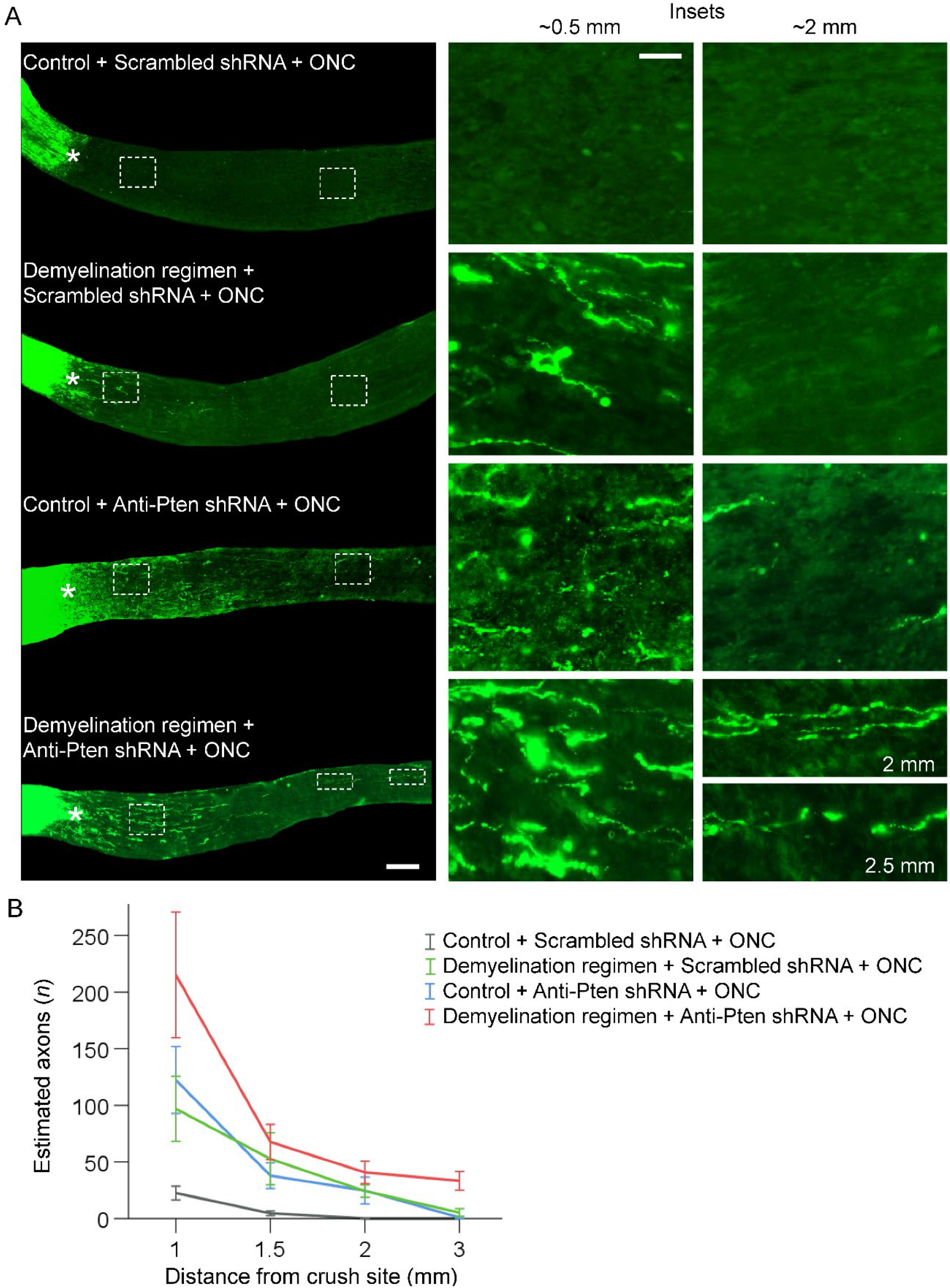
Demyelination regimen in a combination with Pten KD promotes significantly longer axon regeneration than each treatment alone. (**A**) Representative images of the longitudinal optic nerve sections with CTB-labeled axons 2 weeks after ONC and pre-treatment conditions, as marked (experimental timeline in Fig. 1A); **\*** - indicates crush site. Insets: Images of regions proximal and distal to the injury site are magnified for better visualization of regenerating axons or their absence in control. Scale bars, 300 μm (main panels), 100 μm (insets). (**B**) Quantification of regenerating axons visualized by CTB 2 weeks after ONC at increasing distances from the injury site across various conditions, as marked and shown in panel *A* (Mean ± S.E.M; *n* = 5 cases for each condition). ANOVA with repeated measures, sphericity assumed, overall *F* = 8.0, *p* < 0.001; *p*-values of pairwise comparisons by posthoc LSD are shown in Table 2A.

**Fig. 5.**
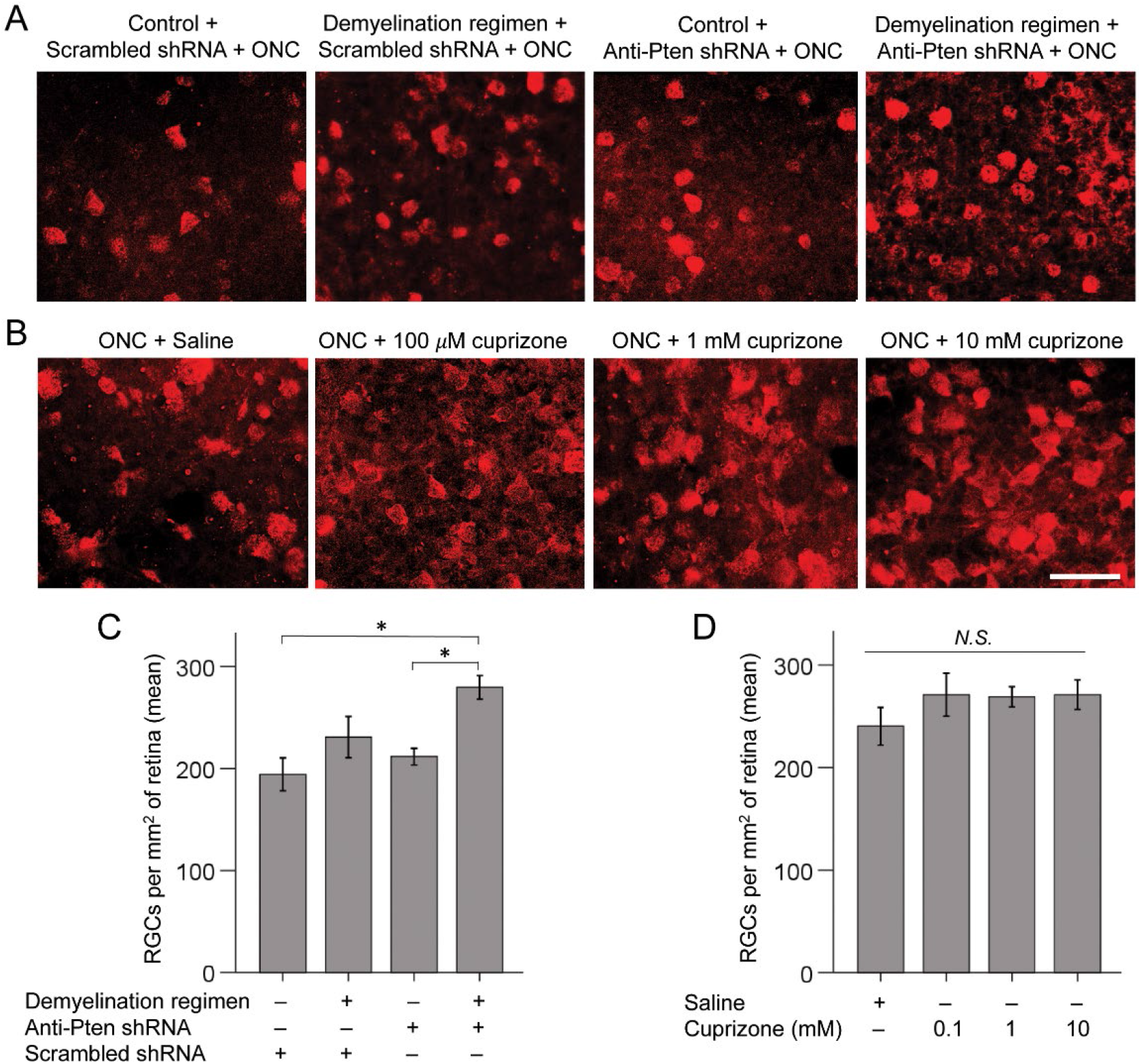
Demyelination regimen in a combination with Pten KD promotes RGC survival. (**A-B**) Representative images of RBPMS-labeled RGCs in flat-mounted retinas 2 weeks after ONC and pre-treatment (*A*) or post-injury treatment (*B*) conditions, as marked (experimental timeline in Fig. 1*A*). Scale bar, 50 μm. (**C-D**) Quantification of RGC survival in retinal flat-mounts immunostained for the RGC marker RBPMS 2 weeks after ONC and pre-treatment (*C*) or post-injury treatment (*D*) conditions, as marked (Mean ± S.E.M shown; *n* = 4-5 cases for each condition). By ANOVA, overall *F* = 3.81 (*C*) and *F* = 0.84 (*D*), *p* < 0.03 (*C*) and *p* < 0.50 (*D*); *p*-values of pairwise comparisons by posthoc LSD are shown in Tables 3A (*C*) and 3B (*D*); * *p* < 0.02; *N*.*S*., not significant. Note: Mice at the time of injury were a month younger for the intravitreal injection of treatments experiment (10 weeks old as typical for this assay), compared to the demyelination regimen experiment, which required prolong pre-treatment with the diet. A slightly higher RGC survival baseline in the intravitreal injection experiment compared to control used in the demyelination regimen experiment is consistent with mice being older in the latter, because older animals typically have more severe deficits after CNS injury.

**Fig. 6.**
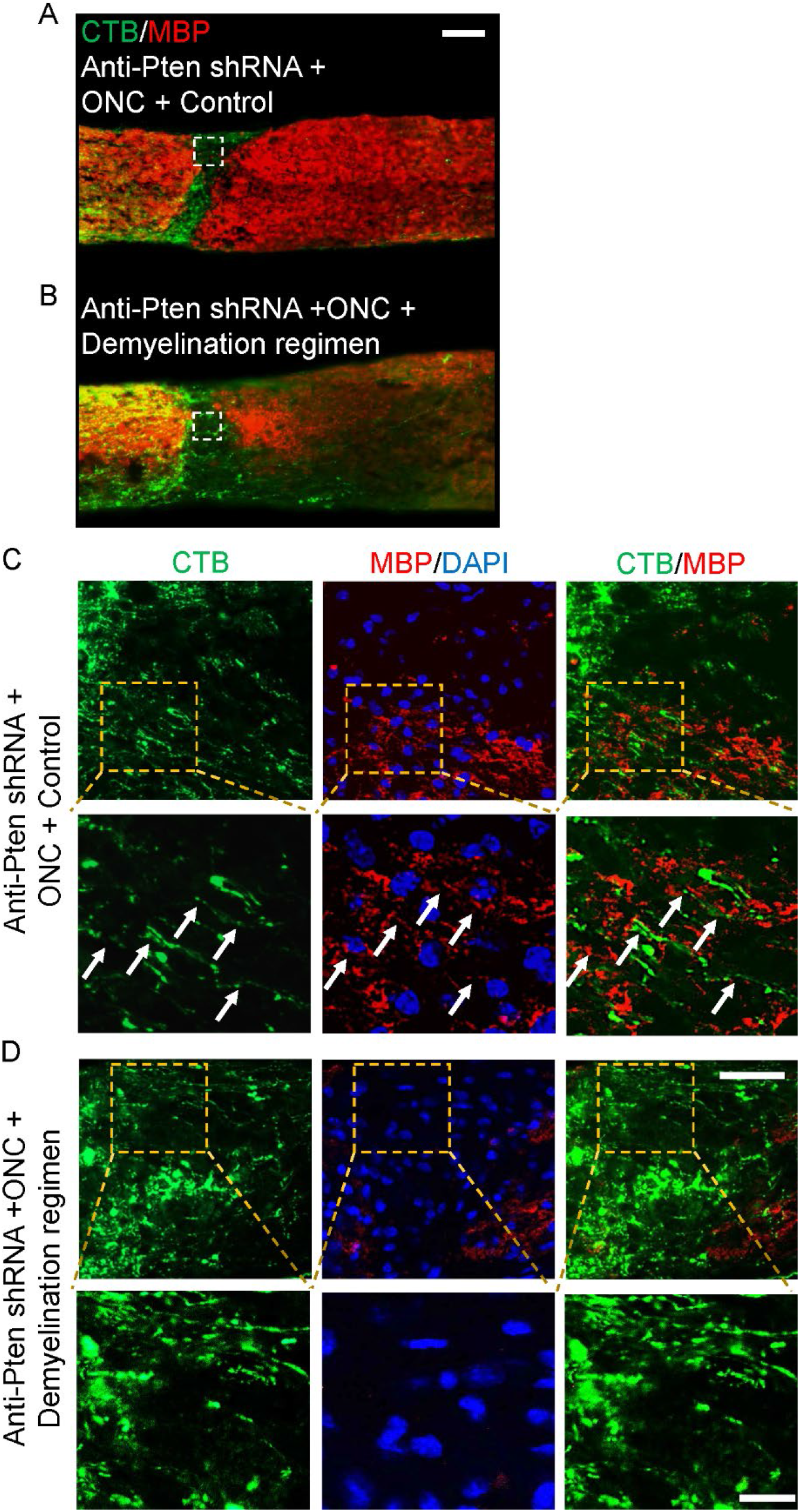
Demyelination regimen prevents premature myelination of the regenerating axons. (**A-B**) Representative images of optic nerve longitudinal sections immunostained for MBP and CTB at 4 weeks after ONC, pre-treated with AAV2 expressing anti-Pten shRNA, and either on normal (*A*) or cuprizone (*B*) diet. Injury site indicated by dashed white lines box. Scale bar, 100 μm. (**C-D**) Confocal images of the injury site regions (outlined in white dashed lines in *A-B*) show Pten KD-stimulated regenerating axons growing through the injury site being myelinated on normal diet (*C*) but unmyelinated on cuprizone diet (*D*). Zoomed-in insets (*lower panels* in *C-D*) of the regions outlined in yellow dashed lines (*upper panels* in *C-D*) show areas containing the regenerating axons visualized by CTB co-localizing with MBP signal (*arrows* in *C*), or the regenerating axons visualized by CTB with no adjacent MBP signal (*D*). Scale bars, 50 μm (main panels), 20 μm (insets).

### Localized treatment with cuprizone promotes axon regeneration

Finally, we asked whether a localized treatment with cuprizone facilitates axon regeneration without rapamycin and without pre-treatment (i.e., cuprizone diet prior to injury). The advantage of the localized treatment is that it enables a higher concentration of cuprizone (that remains within the eye after intravitreal injection and defuses into the adjacent ONC site though the optic nerve head) to target the oligodendrocytes in the injury site without eliciting systemic toxicity. This would not be possible to achieve through the cuprizone diet, as the animals would die rapidly if systemically exposed to such a high concentration of cuprizone. Thus, a localized intravitreal injection of cuprizone was performed immediately after ONC, with 3 μl of 100 μM, 1 mM, and 10 mM solutions. Two weeks after injury, all treatment groups showed a trend towards axon regeneration, but only the 1 mM dose was significant (**Fig. 7A-B** and **Table 2B**). The trend towards a positive effect on RGC survival was not significant (**Fig. 5B,D** and **Table 3B**). Our findings are consistent with prior reports that even untreated RGCs attempt, but fail, to regenerate injured axons^42^, as preventing *de novo* myelination facilitated axonal growth. These findings are also consistent with that RGCs’ intrinsic axon growth capacity declines as they mature^20^, because stimulating the intrinsic mechanism of axon regeneration by Pten KD in a combination with the demyelination regimen led to more robust axon regeneration than either treatment alone.

**Fig. 7.**
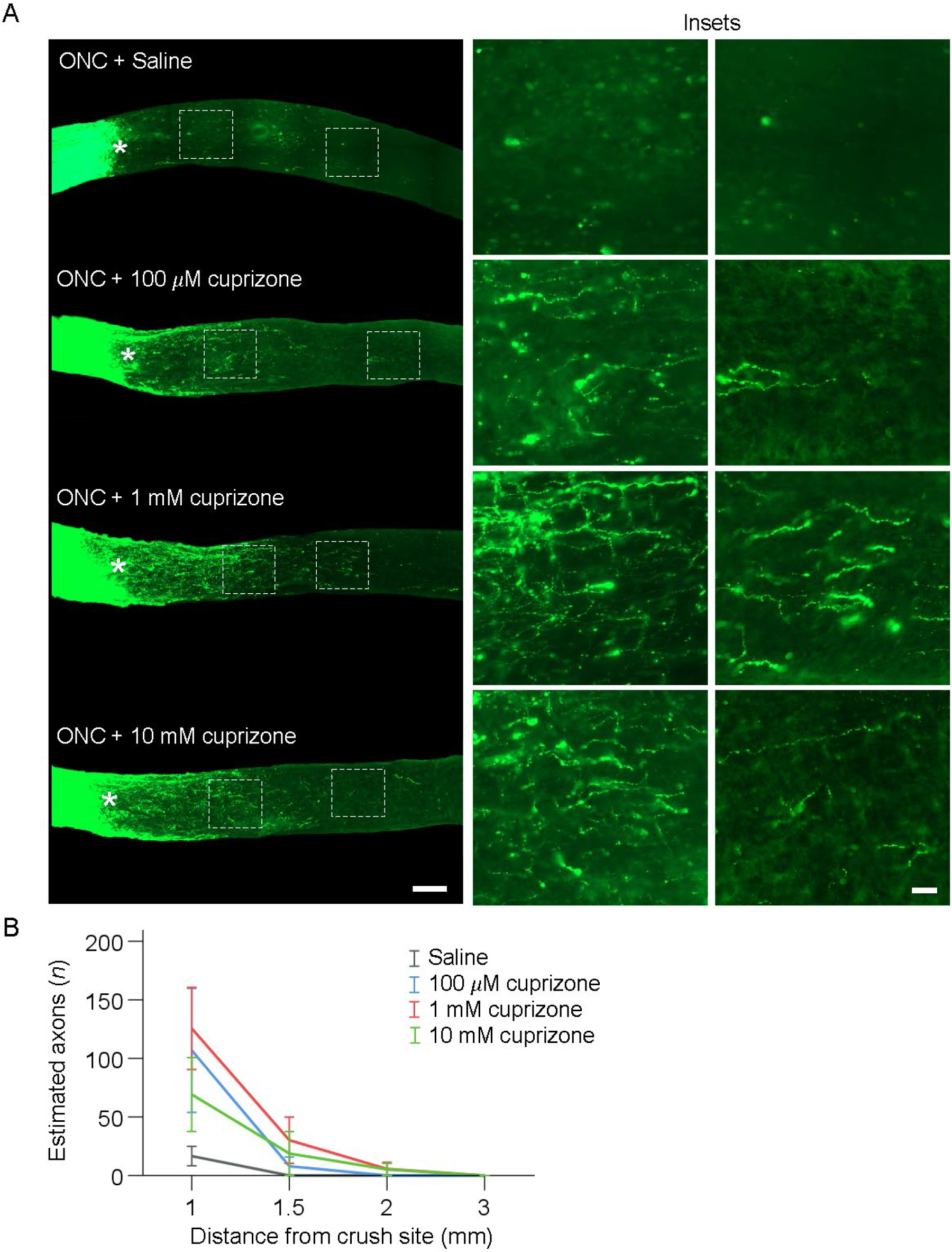
Intravitreal injection of cuprizone after ONC promotes axon regeneration. (**A**) Representative images of longitudinal optic nerve sections showing CTB-labeled axons 2 weeks after ONC, treated immediately after injury with a single intravitreal injection of vehicle or cuprizone at different concentrations, as marked; **\*** - indicates crush site. Insets: Images of regions distal to the injury site are magnified for better visualization of regenerating axons or their absence in control. Scale bars, 300 μm (main panels), 50 μm (insets). (**B**) Quantification of regenerating axons visualized by CTB 2 weeks after ONC at increasing distances from the injury site across various conditions, as marked and shown in panel *A* (Mean ± S.E.M; *n* = 5 cases per condition). ANOVA with repeated measures, sphericity assumed, overall *F* = 2.9, *p* < 0.03; *p*-values of pairwise comparisons by posthoc LSD are shown in Table 2B.

### Cuprizone does not act on RGCs directly or stimulates intraocular inflammation

To test if cuprizone can act on RGCs directly to affect their axon growth, we incubated adult RGCs, isolated by immunopanning for Thy1, for 5 days in culture in a defined growth medium^19^ (see Methods) with varying concentrations of cuprizone. Cuprizone was not toxic to RGCs, but did not promote axon growth either (**Supplementary Figure 8**). To test if cuprizone stimulates intraocular inflammation, which promotes axon regeneration^8^, we examined the retinas for inflammatory fibrotic scarring and immunostained for Ida1 (microglia/macrophage marker) after intravitreal injection of cuprizone (1 mM), zymosan (which promotes axon regeneration through stimulating intraocular inflammation^8^), or PBS. We did found evidence of inflammatory fibrotic scarring in zymosan, but not PBS or cuprizone, treated group (**Fig. 8A**). We also found significant increase in microglia/macrophage number per retina and area per cell (as activated microglia/macrophage change morphologically) in zymosan treated group, but no change on either parameter in cuprizone treated group, compared to the PBS control treated group (**Fig. 8B-D**). This data demonstrates that cuprizone promotes axon regeneration without stimulating inflammation.

**Fig. 8.**
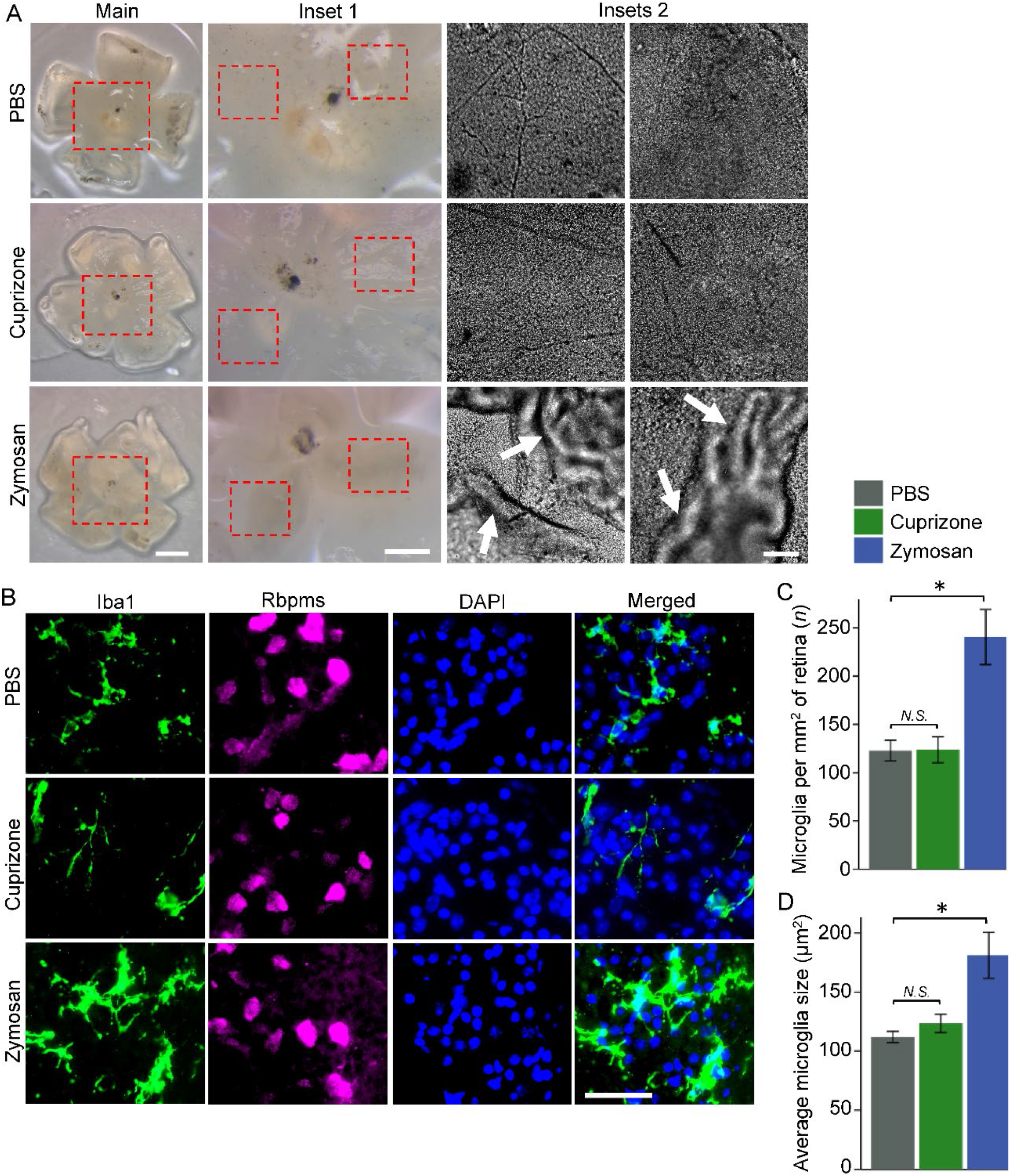
Cuprizone does not stimulate intraocular inflammation. (**A**) Representative images of flat-mount retinas from the eyes treated with PBS (control), 1 mM cuprizone, and zymosan, showing examples of inflammatory fibrotic scarring (indicated by arrows) found in zymosan, but not in PBS or cuprizone, treated eyes. Main panel and inset 1 images were acquired using dissection microscope (Leica M205) with HD digital camera (Leica MC170 HD). Insets 2 images were acquired using fluorescent microscope (Zeiss, AxioObserver.Z1). Scale bars: main, 1 mm; inset 1, 500 μm; insets 2, 100 μm. (**B**) Representative images of flat-retinal thin sections (containing the ganglion cell layer) immunostained for Iba1 (microglia/macrophage marker), Rbpms (RGC marker), and DAPI (nuclear marker) at 2 weeks after ONC and intravitreal treatment with PBS (control), 1 mM cuprizone, and zymosan. Images were acquired using fluorescent microscope (Zeiss, AxioObserver.Z1). Scale bars: 50 μm. (**C-D**) The difference between number of microglia per retina or microglia size (visualized by immunostaining for the Iba1 marker of microglia/macrophage) was not significantly different between PBS and cuprizone treated eyes, but was significant between PBS and zymosan treated eyes, by ANOVA with posthoc LSD for pairwise comparisons (* *p* < 0.01 in *C* and *D* by posthoc; overall *F* = 14.4 and *p* < 0.01 in *C*, and overall *F* = 8.9 and *p* < 0.02 in *D*). *N* = 3, *n* < 40 (at least 40 cells quantified per case); error bars = SEM.

### Post-injury-born oligodendrocytes express axon growth-inhibitory membrane proteins

Myelin debris-associated inhibitors of axon growth also include membrane proteins, which are presented on the surface of live oligodendrocytes to the axonal stamp or growth cone as the axons attempt to regenerate after optic nerve injury, without needing to be exposed to axonal receptors specifically on myelin debris only after oligodendrocyte’s death. Such myelin-associated inhibitors that are presented on the surface of oligodendrocytes include, NogoA^68^ and Omgp^69^. BAIs also were recently discovered to inhibit axon growth by live oligodendrocytes co-cultured with neurons^27^. Thus, we investigated whether these inhibitors of axon growth are expressed by post-injury-born immature oligodendrocytes. We found that NogoA (Rtn4) was expressed in all stages of oligodendrocytes maturation (see Fig. 2F), including in post-injury-born NFOs. Rtn4 expression levels were significantly higher in the NFOs and in the MOs (but not in the sSOs) from the injured compared to uninjured conditions, and within the injured condition expression levels were significantly higher in the MOs (but not in the sSOs) compared to NFOs (**Fig. 9A**). Omgp (Omg) was expressed, but at a lesser level in the injured compared to uninjured conditions in all stages of oligodendrocytes maturation, and within the injured condition expression levels were higher in the MOs (but not in the sSOs) compared to NFOs (**Fig. 9B**). BAI1 (Adgrb1) was expressed significantly higher in the NFOs, and its expression levels were similar for each stage between uninjured and injured conditions (**Fig. 9C**). BAI2 (Adgrb2) also was expressed primarily in the NFOs, but its levels of expression were significantly higher in the NFOs and in intermediate stage of maturing oligodendrocytes in the injured compared to uninjured conditions (**Fig. 9D**). BAI3 (Adgrb3) levels of expression bordered noise at all stages of oligodendrocytes maturation in both conditions. These data suggest that, axon regeneration-inhibitory NogoA and BAI1-2 are moderately expressed in post-injury-born immature oligodendrocytes, whereas Omgp is expressed at a lesser level in the injured condition, and BAI3 expression is bordering noise in either condition. Thus, axon growth-inhibitory membrane proteins, NogoA, Omgp, and BAI1-2, are expressed by live immature post-injury-born oligodendrocytes, positioned to stall axonal regeneration during premature myelination.

**Fig 9.**
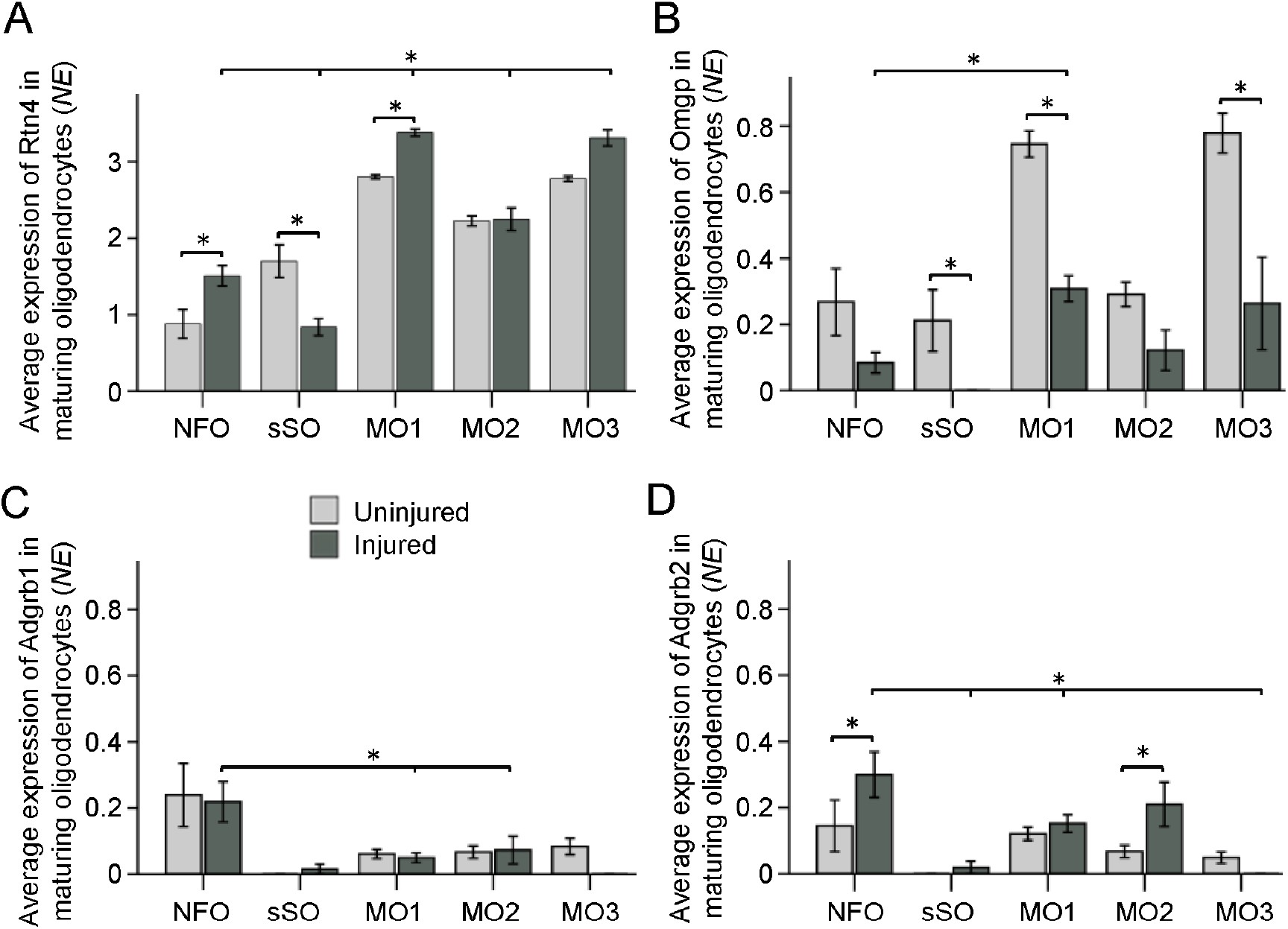
Post-injury-born oligodendrocytes express genes encoding axon growth-inhibitory membrane proteins. (**A**) NogoA (Rtn4) expression levels significantly higher in the NFOs and in MO1 (but lower in the sSOs) from the injured compared to uninjured conditions, and within the injured condition expression levels significantly higher in the MO1-3 (but lower in the sSOs) compared to NFOs (\**p* < 0.01). (**B**) Omgp expressed at a significantly higher levels in MO1 compared to NFOs within the injured condition (\**p* < 0.03), but expressed significantly less in the injured compared to uninjured sSOs, MO1, and MO3 (\**p* < 0.05). (**C**) BAI1 (Adgrb1) expressed significantly higher in the NFOs compared to sSOs, MO1, and MO2 within the injured condition (\**p* < 0.03). (**D**) BAI2 (Adgrb2) expression significantly higher in the NFOs and MO2 from the injured compared to uninjured conditions (\**p* < 0.03), and within the injured condition expression is significantly higher in the NFOs compared to sSOs, MO1, and MO3 (\**p* < 0.03). Significant differences (*p-*values) determined by 2-way ANOVA with posthoc LSD pairwise comparisons. Error bars = SEM. *NE =* normalized expression.

## DISCUSSION

A major barrier in experimental axon regeneration research is that CNS neurons that respond to regenerative treatments stall axon growth prior to reaching post-synaptic targets (e.g.,^2,13–15,17–21^). Our data shows that post-injury-born immature oligodendrocytes integrate into the glial scar, and that preventing oligodendrocytes from prematurely myelinating the growing axons facilitates axon regeneration.

We also found that, ONC kills most oligodendrocytes in the injury site and their myelin debris fill the injury site (as we observed at 3 days after ONC), however, by 2 weeks after ONC myelin debris are cleared from the injury site and newly-born oligodendrocytes fill (more than in uninjured tissue) the injury site. Thus, while myelin debris from dead oligodendrocytes are cleared overtime, newly-born oligodendrocytes become a major cellular component of the injury site glial scar, positioning them to inhibit the axons that attempt to regenerate. We show that cuprizone treatment decreases newly-born oligodendrocytes in the injury site, thereby promoting axonal regeneration. This finding provides an important insight into why axons stall regeneration, and prompts future studies to investigate whether therapeutically suspending myelinating activity throughout the optic nerve and tract (and not only at/near the glial scar) would facilitate full-length experimental axon regeneration.

An alternative possibility that we have addressed is whether cuprizone may act on RGCs directly to stimulate their axonal growth. As we found no increase in axon growth of adult RGCs incubated with various doses of cuprizone even after 5 days in culture, we concluded that cuprizone does not stimulate axon regeneration by acting on RGCs directly. However, even at the higher, 10 mM, concertation, cuprizone was still not toxic to RGCs in culture, consistent with its established preferential toxicity for oligodendrocytes and not neurons. Another alternative possibility that we have addressed is whether cuprizone may promote axon regeneration through stimulating inflammation^8^. As we found that intravitreal injection of cuprizone neither activated the marker (Iba1) of intraocular inflammation nor caused fibrotic retinal scarring, in contrast to zymosan which activated Iba1 and caused fibrotic retinal scarring, we concluded that cuprizone promotes axon regeneration without stimulating inflammation.

While cuprizone does not affect astrocytes directly^29^, it can indirectly lead to their activation^70^ and secretion of CSPG in the glial scar. Co-concurrent targeting of CSPG^71^ and intravitreal injection of cuprizone, along with Pten or Klf9 KD in RGCs^15^ may lead to more robust axon regeneration than each on its own. After cuprizone treatment is discontinued^32,33^, new and recovered oligodendrocytes can myelinate the regenerated axons that reached their respective postsynaptic targets; however, even the reversible side effects of such a cuprizone treatment may be overly concerning for clinical use. Nevertheless, our findings point to a new direction for repairing injured CNS axonal connections, which in the future may involve localized, more efficient, and less toxic treatments that will temporarily suspend active myelination by oligodendrocytes.

As a part of these studies, we also generated a transcriptomic resource for studying gene networks and pathways in the optic nerve OPC and oligodendrocyte subtypes under physiological and injured-pathophysiological conditions. We thus presented a Subtype Gene Browser website (in the same format as we previously did for the RGC subtypes^61^), which provides a platform for comparing the gene expression in the resource-dataset we generated. This tool will assist the scientific community in the investigation of the molecular and physiological differences between optic nerve OPC and oligodendrocyte subtypes under physiological and injured-pathophysiological conditions.

Through which mechanism does myelination of the regenerating axons inhibit their growth? Perhaps, active myelination provides the myelin-associated inhibitors in closer proximity and across a larger axonal area than would myelin debris. This increase in inhibitory molecules may reach a threshold of intracellular signaling that outcompetes the pro-axon-growth program. Some of the prominent myelin-associated inhibitors of axon growth, NogoA^68^ and Omgp^69^, are membrane proteins that can be presented by live immature oligodendrocytes to inhibit axon regeneration, without needing to be exposed to axonal receptors specifically on myelin debris only after oligodendrocyte’s death. BAIs also were recently discovered to inhibit axon growth by live oligodendrocytes cocultured with neurons^27^. Indeed, we found that NogoA, Omgp, and BAI1-2 are moderately expressed in oligodendrocytes newly-born after optic nerve injury, through which they could inhibit axon regeneration. Alternatively, the axon-wrapping process/sheath may express certain ligands only during the active wrapping period, which signal the axon to stop growing. It is also possible that during the growth state axons require more nutrients, which are less accessible after they are wrapped in myelin sheaths.

Importantly, recent studies argued for therapeutically promoting myelination of the experimentally-stimulated to regenerate at early stage of growth, before the axons reached their post-synaptic targets^72^. Our study, however, suggests that myelinating activity should be therapeutically suspended until the axons regenerate the full-length to reach their targets. Thus, considering that cuprizone only modestly diminishes optic nerve oligodendrocytes distal from the injury site, would conditional genetic suspension of myelinating activity within the optic nerve and tract after ONC facilitate full-length experimental axon regeneration?

Recent studies have shown that RGC subtypes differ in responsiveness to experimental axon-regeneration treatments^61,73–75^, and it is possible that the regenerating axons of certain RGC subtypes become susceptible to myelination later than others^76^ and therefore stall growth later. In contrast to normal developmental axon growth, where myelination in the optic nerve begins over a week after the axons have reached their targets in the brain^23,24^, oligodendrocytes in the adult optic nerve, which are either surrounding the injury site and beyond, or are newly-formed post-injury from the optic nerve OPCs, are readily available to myelinate the regenerating axons while they are still growing (prior to reaching synaptic targets). The embryonic growing axon diameter is thin (compared to a fully mature axon), and it is only after they stop growing and form synapses that their dimeter increases^77^. The onset of myelination is triggered by an increase in axonal dimeter to a certain threshold^78^, and by the time optic nerve matures all the axons of varying diameters are myelinated^79^. We therefore hypothesized that the embryonic cell state would facilitate the regeneration of thin axons, as all axons during embryonic development are thin while still in the axonal growth stage. Indeed, we used single cell transcriptome profiling to analyze the RGCs which regenerated axons in response to Pten KD, and found that the adult RGCs which regenerated axons in response to Pten KD shifted transcriptomically (based on the transcriptome similarity) towards an embryonic RGC state^80^. Thus, it would be important for future studies to further investigate the hypothesis raised by our studies, which suggests that the experimental treatments need to aim at reverting the RGCs towards an embryonic state, when their intrinsic capacity for axon growth is higher^19,20,81,82^ and susceptibility to myelination is lower^80^.

## METHODS

### Animal Use, Diet, and Surgeries

All experimental procedures were performed with the approval of the Institutional Animal Care and Use Committee and of the Institutional Biosafety Committee at the University of Connecticut Health Center and performed in accordance with the ARVO Statement for the Use of Animals in Ophthalmic and Visual Research. Mice were housed in the animal facility with a 12-h light/12-h dark cycle (lighting on from 7:00 AM to 7:00 PM) with a maximum of five mice per cage. Wild-type 129S1/SvlmJ male mice were obtained from the Jackson Laboratory. Food and water were available *ad libitum*. To induce demyelination in the CNS, starting at 8 weeks of age, 0.3% cuprizone [bis(cyclohexanone)oxaldihydrazone] (MilliporeSigma) was mixed with normal chow (containing crushed Teklad global 19% protein rodent diet; Envigo), and mice started receiving, 5 days per week, intraperitoneal injections of 10mg/kg rapamycin (LC Laboratories) dissolved in a vehicle solution of 5% ethanol in phosphate-buffered saline (PBS; Fisher Scientific), as described^32,33^. Cuprizone was decreased to 0.2% at week 4 of the diet and then to 0.1% at week 8 of the diet. Normal chow diet and vehicle intraperitoneal injections were used for control in age-matched mice. Rapamycin and control injections were discontinued after 6 weeks, when viruses with anti-Pten or scrambled shRNAs were injected, because rapamycin inhibits axon regeneration^34^. Giving a 2-week period for clearance of rapamycin before ONC is on the stringent side, because rapamycin’s half-life after injection is under 3 days, with complete clearance of even higher doses (than we used) from the organism in just several days^35,36^. To stimulate axon regeneration, 3 μl of adeno-associated virus serotype 2 (AAV2, 1 × 10^12^ GC/mL; Cyagen Biosciences, Inc.) expressing anti-Pten (target sequences: 5′-GCAGAAACAAAAGGAGATATCA-3′, 5′-GATGATGTTTGAAACTATTCCA-3′, 5′-GTAGAGTTCTTCCACAAACAGA-3′, and 5′-GATGAAGATCAGCATTCACAAA-3′) or scrambled control (sequences: 5’-TCGAGGGCGACTTAACCTTAGG-3’) shRNAs were injected intravitreally (avoiding injury to the lens) 2 weeks prior to ONC. AAV2 is typically used in this model, because AAV2 (but not other AAV serotypes) preferentially transduces RGCs when injected locally into the eye vitreous. Untreated, RGCs die progressively over time after ONC. Thus, a standard protocol in the field is to inject AAV2 two weeks prior to ONC, which allows sufficient time for transduction and target gene KD^15,18^ (the timing of treatments and injury are shown in Figure 1A). Because remyelination becomes notable only in about a week after discontinuation of the cuprizone diet^32,33^, cuprizone was excluded from the diet one week prior to sacrifice at 2 weeks after ONC (week 9 of the diet), except when mice were sacrificed at 4 weeks after ONC (in which case, cuprizone diet was started 5 weeks before ONC and was stopped one day before sacrifice). Intravitreal injections of zymosan (3 μl; Z4250, MilliporeSigma), or cuprizone dissolved in saline, were performed right after ONC surgery in 10-week old mice: 3 μl of 100 μM, 1 mM, or 10 mM cuprizone solutions were injected; vehicle was used for control injections. Cholera toxin subunit beta (CTB, 1% in 2 μl PBS; 103B, List Biological) axonal tracer was injected intravitreally one day prior to sacrifice. Optic nerve surgeries and intravitreal injections were performed under general anesthesia, as described^2,15^. The animals were euthanized 2, 4, or 6 weeks after ONC for histological analysis.

### Histological Procedures

Standard histological procedures were used, as described previously^2,15^. Briefly, mice were anesthetized and transcardially perfused with 0.9% saline solution followed by 4% paraformaldehyde (PFA) at 3 days, or 2, 4, or 6 weeks, after optic nerve injury. The optic nerves, eyes, and brains were resected. The eyes were post-fixed 2 hours at room temperature and the optic nerves and brains for 24 hours at 4 °C. The optic nerves and brains were then transferred to 30% sucrose overnight at 4 °C. The retinas (for analyzing RGC survival and fibrotic scar) were resected from the eyes and kept in PBS at 4 °C. The optic nerves, flattened retinas (for analyzing microglia), and brains were embedded in OCT Tissue Tek Medium (Sakura Finetek), frozen, and cryostat-sectioned at 14 μm longitudinally for the optic nerves, horizontality (capturing the ganglion cell layer of the whole retina) for flattened retinas, and at 20 μm coronally for the brains. The optic nerve cryosections were immunostained on Superfrost Plus glass slides (VWR International, LLC), and the brain cryosections and the retinas were immunostained, free-floating in wells (24-well plate), and then transferred onto the coated glass slides after making 4 symmetrical slits in the retinas for flat-mounting, and then mounted for imaging. For immunostaining, the tissues were blocked with appropriate sera, incubated overnight at 4 °C with primary antibodies as indicated, then washed three times, incubated with appropriate fluorescent dye-conjugated secondary antibodies (1:500; Alexa Fluor, Life Technologies) overnight at 4 °C, and washed three times again.

### Imaging and Quantifications of Histological Tissue Preparations

CC1, MBP, GFAP, and Aspa signals in the optic nerves and brains were visualized by immunostaining, using antibodies for CC1 (1:500; mouse IgG2b, ab16794, Abcam), MBP (1:500; rabbit, ab40390, Abcam), GFAP (1:400; rabbit, ab7260, Abcam), Aspa (1:500; rabbit, ab223269, Abcam), and counterstained with DAPI (1:5000; Life Technologies) to label nuclei. *Z*-stacked images of the optic nerves, with 5 planes at 0.5 μm intervals, were acquired using a fluorescent microscope (40x/1.2 C-Apochromat W; AxioObserver.Z1, Zeiss) and merged (ZEN software, Zeiss). Note: Although 2 hours post-fix at room temperature is common for the optic nerves and is appropriate for most antibodies, for visualizing CC1 signal (using the antibody above) in the ONC site, it is necessary to post-fix for 24 hours at 4 °C. For quantification of MBP and CC1 signal in the optic nerves, average fluorescence signal intensity along the optic nerve diameter at 50 μm intervals throughout 250 μm distance spanning the injury site, or in an equivalent region of an uninjured optic nerve, was measured using the ImageJ plugin Plot Profile. Averages of signals from 3 longitudinal tissue sections per optic nerve were quantified for each biological replicate (*n*) per condition. Individual planes from the *z*-stacks were used for representative images. MBP and CC1 signals in the corpus callosum were visualized similarly, and average fluorescence signal intensity was measured using the ImageJ Measure tool in the middle of corpus callosum (344 μm x 149 μm area) in 3 coronal tissue sections per brain, collected between Bregma-1.58 mm ~ −0.94 mm coordinates. Averages of signals from 3 tissue sections were quantified for each biological replicate (*n*) per condition. To quantify regenerated axons in the optic nerve, axons were visualized at 2 weeks after optic nerve injury by immunostaining with the anti-CTB antibody (1:500; rabbit, GWB-7B96E4, GenWay) and fluorescent dye-conjugated secondary antibodies (1:500; Alexa Fluor, Life Technologies); 2 μl of CTB (1% in PBS) was injected intravitreally 1 day prior to sacrifice. No spared axons were found in controls or experimental conditions (i.e., at 2 weeks after injury, no axons were found at the region of the optic nerve most distal to the injury, where regenerating axons have not yet reached at this time point)^2,15^. Regenerated axons (defined as fibers continuous for >100 μm, which are absent in controls and are discernible from background puncta and artifactual structures), were counted manually using a fluorescent microscope (40x/1.2 C-Apochromat W; AxioObserver.Z1, Zeiss) in at least 3 longitudinal tissue sections per optic nerve at 0.5 mm, 1 mm, 1.5 mm, 2 mm, and 3 mm distances from the injury site (identified by the abrupt disruption of the densely packed axons near the optic nerve head, as marked by an asterisk in the figures), and these values were used to estimate the total number of regenerating axons for each optic nerve (biological replicate, *n*) per condition, as described^13,15,21^. For representative images, serial fields of view along the longitudinal optic nerve tissue section were imaged as above; *z*-stacks with 5 planes at 0.5 μm intervals were deconvoluted, merged, and stitched. Then, processed images of 3 tissue sections from the same optic nerve were superimposed over each other and merged using Photoshop CS6 (Adobe), shown as representative images. To visualize co-localization of MBP and CTB signals in the optic nerve 4 and 6 weeks after ONC, and to visualize segregation between CC1 and GFAP or Aspa signals in the optic nerve 2 weeks after ONC, z-stacks with 0.5 μm intervals were acquired using confocal microscopy (40x/1.3 Achrostigmat Oil; LSM 880, Zeiss). For quantification of RGC survival, flatmounted retinas were immunostained with an antibody for the RGC-specific marker, RBPMS^67^ (1:500; guinea pig, 1832-RBPMS, PhospoSolutions), and RMPBS+ cells were counted as described^13,15,21^ using ImageJ software. RGCs were counted in at least 4 images per retina, acquired at prespecified areas using a fluorescent microscope (40x/1.2 C-Apochromat W; AxioObserver.Z1, Zeiss) as above, approximately at 2 mm from the optic nerve head in four directions within the ganglion cell layer per retina, then averaged to estimate overall RGC survival per mm^2^. For visualization of fibrotic scarring, flat-mounted retinas were imaged using microscope (Leica M205) with HD digital camera (Leica MC170 HD), and insets were imaged using microscope (Zeiss, AxioObserver.Z1). For quantification of microglia/macrophage, thin cryosections of the flattened retinas (containing the ganglion cell layer) were immunostained for RBPMS (as above) and for microglia/macrophage-specific marker, Iba1 (1:600; rabbit, 019-19741, Wako). Then, images were acquired at prespecified areas using a fluorescent microscope (40x/1.2 C-Apochromat W; AxioObserver.Z1, Zeiss) and quantified as above using ImageJ software, to estimate overall average microglia/macrophage number per mm^2^. To measure microglia/macrophage size, Iba1+ area for individual microglia/macrophages was quantified in μm^2^, using Analyze Particles function of ImageJ software, and averaged per retina. Pten KD in RGCs by anti-Pten shRNA was validated as previously described^18^. Briefly, flat-mounted retinas transduced either with anti-Pten shRNA AAV2 or scrambled shRNA AAV2 were immunostained, 2 weeks after transduction, with an antibody for Pten (1:200; rabbit, 9559, Cell Signaling Technology), neuronal marker βIII-Tubulin (1:500; mouse IgG2a, MMS-435P, BioLegend), and counterstained with DAPI (1:5000; Life Technologies) to label nuclei. *Z*-stack images of the ganglion cell layer were acquired using confocal microscopy as described above, and orthogonal projections through Z-stock used for representative images.

### RGC Culture and Immunostaining

RGCs were purified from both sexes of 5 10-week old adult mice retinal single cell suspension by immunopanning for Thy1 (CD90, MCA02R, Serotec) after depletion of macrophages (using anti-mouse macrophage antibody, AIA31240, Accurate Chemical) and washing off the nonadherent cells. RGCs were plated and cultured in defined growth medium following our established protocol^19,83,84^, with the following modifications for adult tissue: digestion in papain decreased to 10 minutes, amount of papain increased by 25%, only one macrophage depletion plate used for 20 minutes, Thy1 panning dish shaken more stringently, culture plates coated with a modified growth substrate, and growth factors and sato supplement doubled in the defined growth medium. RGCs were incubated with varying concentrations (as indicated in Fig. 7) of cuprizone (MilliporeSigma) or without it. After 5 days in culture, RGCs were fixed and immunostained with neuronal marker βIII-Tubulin (1:500; mouse IgG2a, MMS-435P, BioLegend) and DAPI (1:5000; Thermo Fisher Scientific). AlexaFluor fluorophore-conjugated secondary antibody (1:500; Thermo Fisher Scientific) were used for fluorescent microscopy (with AxioObserver.Z1, Zeiss).

### Statistical Analyses

All tissue processing, quantification, and data analyses were done masked throughout the study. The animals on demyelination or normal diet were randomly selected from within each group for experimental or control AAV2 injections, and the investigators performing the surgeries and quantifications were masked to the treatments’ identities. Sample sizes were based on accepted standards in the literature and our prior experiences^2,15^. Sample size (*n*) represents a total number of biological replicates in each condition. All experiments included appropriate controls. No cases were excluded in our data analysis, although a few animals that developed a cataract in the injured eye were excluded, and their tissues were not processed, and a few animals died. The data were analyzed by one or two way ANOVA with or without repeated measures, as appropriate, and a posthoc LSD for pairwise comparisons, or independent samples *t*-test, as indicated (SPSS). Fold change significance in Fig. 2E was determined using the EdgeR algorithm^85^.

### Single cell RNA-seq (scRNA-seq)

scRNA-seq was performed using the methods we described previously^61^, with modifications described below. Briefly, cells were purified from uninjured (*n* = 5) and injured (*n* = 5) optic nerves of both sexes 10-week old mice. ONC injury was performed at 8 weeks, as described above. Single cell suspension was prepared using slightly modified version of the method described above for obtaining single cell retinal suspension (in order to immunopan the RGCs). Cells were resuspended in DPBS with 0.04% BSA, and immediately processed as follows. Cell count and viability were determined using trypan blue on a Countess FL II, and 6,000 cells from each sample were loaded in parallel for capture onto the Chromium System using the v2 single cell reagent kit (10X Genomics). Following capture and lysis, cDNA was synthesized and amplified (12 cycles) as per manufacturer’s protocol (10X Genomics). The amplified cDNA from each channel of the Chromium System was used to construct an Illumina sequencing library and sequenced on HiSeq 4000 with 150 cycle sequencing. Illumina basecall files (*.bcl) were converted to FASTQs using CellRanger v1.3, which uses bcl2fastq v2.17.1.14. FASTQ files were then aligned to mm10 mouse reference genome and transcriptome using the CellRanger v1.3 software pipeline with default parameters (as described^86^), which demultiplexes the samples and generates a gene versus cell expression matrix based on the barcodes and assigned unique molecular identifiers (UMIs) that enable determining the individual cells from which each RNA molecule originated.

For determining gene expression, normalization of the raw counts was performed using Seurat (v. 4.0.3^87,88^) function, which divides the feature counts by the number of counts per each cell and then applies natural log transformation, resulting in normalized expression (*NE*) values^87,88^. Dimensionality reduction and 2D visualizations were performed using the uniform manifold approximation and projection (UMAP) implementation^89,90^ in Seurat v. 4.0.3 using default parameters^87,88^. 3D visualization and pseudotimeline trajectory were obtained using Monocle3 v. 0.2.3.0 software with default parameters. Clustering of cells comprising oligodendrocyte lineage into subtypes was also preformed using Seurat function with default parameters. For clustering, principal components of gene expression across cells were determined using the top 2000 most variable genes, selected by Seurat’s FindVariableGenes function using the “vst” method. Sex-specific genes (e.g., male Eif2s3y and Ddx3y, and female Xist), cell cycle genes, and mitochondrial genes were regressed out during the scaling of the data but were retained in the dataset for downstream analyses after clustering. A total of 2,496 cells (equal number of cells from injured and uninjured optic nerves after subsampling) that passed quality control (QC) were used. QC filters/thresholds included the following criteria per cell: a maximum threshold of 20% mitochondrial genes expressed in the transcriptome, a minimum of 200 genes, and a maximum of 150,000 UMIs (to mitigate the presence of cell doublets). Gene markers used to identify cell types (see Supplementary Figure 3): CD45, Ida1, and C1qc for immune cells^44–46^, MAG, MBP, and Aspa for oligodendrocytes^50,55^, Pdgfra for OPCs^53^, Fyn for NFOs^55^, GFAP for astrocytes^55^, Col1a1 and Igfbp6 for pericytes/fibroblasts^55^, and Vwf for endothelial cells^91^.

### Design of the website and online tools

The Subtypes Gene Browser was designed in the same format as we previously did for RGC subtypes^61^, using R and ShinyApps with R-markdown language^64^. Boxplots, violin plots, and bar plots were adapted from ggplot2 R software package for data visualization^62^.

### Data Availability

The scRNA-seq data from OPCs and oligodendrocytes that we generated for these studies is available through the NCBI GEO accession *Pending*. Access to the dataset is also available through a user-friendly Subtypes Gene Browser web application, https://health.uconn.edu/neuroregeneration-lab/subtypes-gene-browser.

## ACKNOWLEDGMENTS

This work was supported by grants from The University of Connecticut School of Medicine, Start-Up Funds (to E.F.T.), the Connecticut Institute for the Brain and Cognitive Sciences (Research Seed Grant, to E.F.T. and S.J.C.), the BrightFocus Foundation (Grant G2017204, to E.F.T.), and the National Institutes of Health (NIH) (Grant R01-EY029739, to E.F.T.). Portions of this research were conducted at the High Performance Computing Facility, University of Connecticut. We are grateful to Paul Robson and Greg Sjogren (The Jackson Laboratory for Genomic Medicine, Farmington, CT) for Single Cell RNA-Sequencing service, Sophan Iv and Vijender Singh (Research IT Services, University of Connecticut), and Stephen King (High Performance Computing Facility, University of Connecticut) for assistance with the development of the website and online tools. We thank the University of Connecticut School of Medicine faculty and students, Stephen Crocker, PhD, Robert Clark, MD for advice, Robert Tocci, Dan Thadeio, Nicholas Wasko, Alexela Hoyt, Emmalyn Lecky, Tyler Steidl, Mahit Gupta, and Anthony Antony (undergraduate and graduate students, University of Connecticut) for technical assistance.

## AUTHOR CONTRIBUTIONS

J.X., B.A.R., J.K., and A.L performed the experiments. M.S.S. and A.D. assisted with performing the experiments.

E.F.T. designed the study and wrote the manuscript.

