## Supplemental Information for "Post-injury born oligodendrocytes integrate into the glial scar and inhibit growth of regenerating axons by premature myelination"

Jian Xing et al.

### Supplementary Information

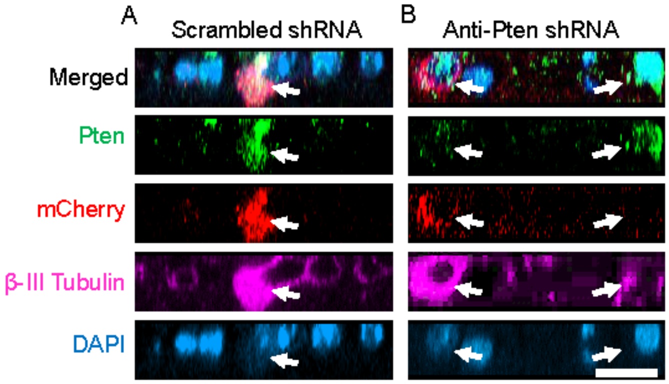


**Supplementary Figure 1. Validation of Pten KD in RGCs.** (**A**-**B**) Flat-mounted retinas transduced either with scrambled shRNA AAV2 (*A*) or anti-Pten shRNA AAV2 (*B*) co-expressing mCherry reporter, immunostained with an antibody for Pten, neuronal marker βIII-Tubulin, and counterstained with DAPI to label nuclei. Representative images show orthogonal projections through confocal *z*-stack images of the ganglion cell layer. In *A*, arrow indicates a control transduced (mCherry) RGC (βIII-Tubulin) robustly expressing Pten. In *B*, arrow on the left indicates control transduced (mCherry reporter) RGC (βIII-Tubulin) not expressing Pten, whereas the arrow on the right indicates a non-transduced (no mCherry) RGC (βIII-Tubulin) robustly expressing Pten. Scale bar, 20 μm.

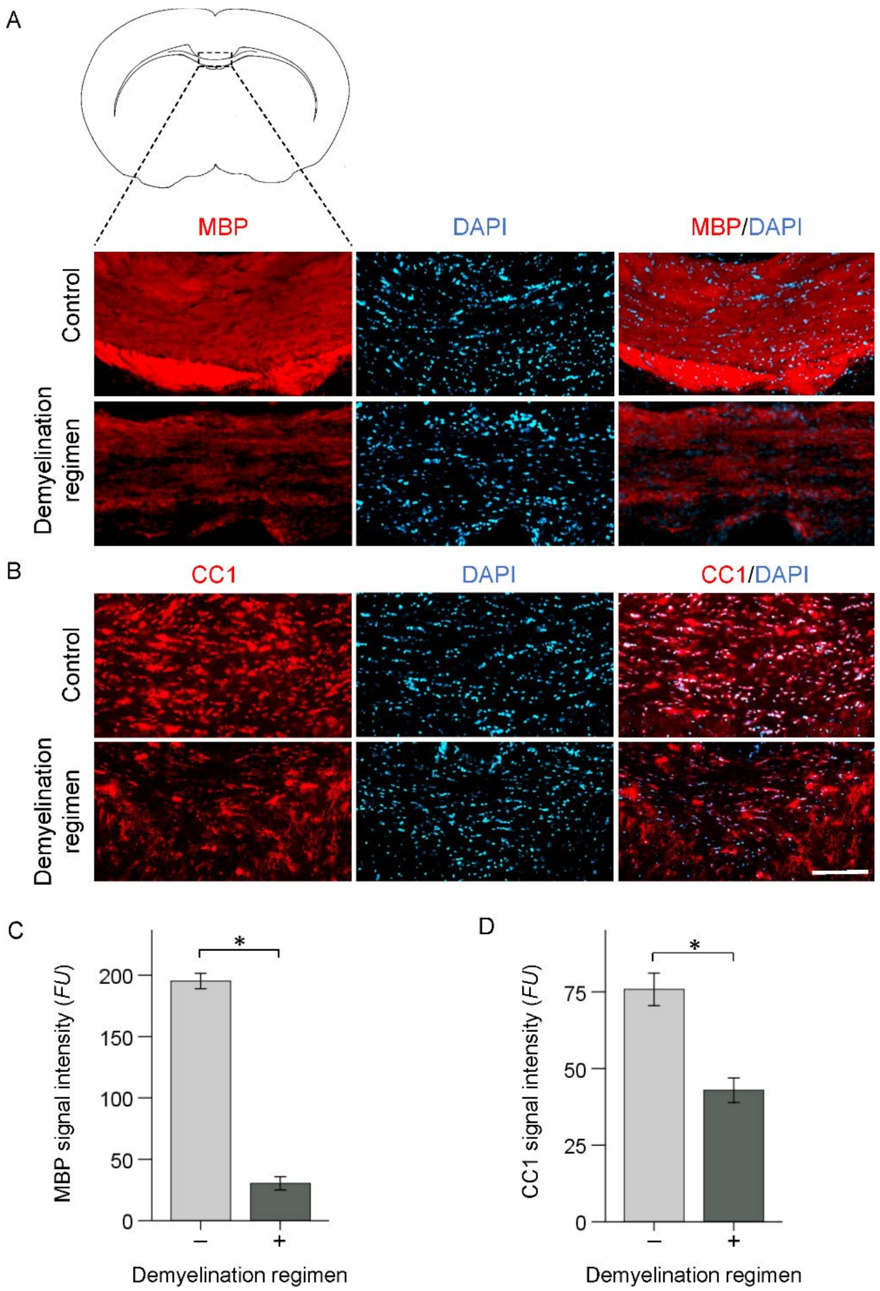


**Supplementary Figure 2. Loss of myelin basic protein (MBP) and mature oligodendrocyte marker CC1 in the corpus callosum after demyelination regimen.** (**A**) Representative coronal images of the corpus callosum immunostained for MBP without (upper panel) or with (lower panel) demyelination regimen. Scale bar, 100 μm. (**B**) Representative coronal images of the corpus callosum immunostained for oligodendrocyte marker CC1 without (upper panel) or with (lower panel) demyelination regimen. Scale bar, 100 μm. (**C-D**) Quantification of MBP (*C*) and CC1 (*D*) immunofluorescence signal intensity, represented in fluorescent units (*FU*s), in the corpus callosum, as marked (Mean ± S.E.M shown; $n = 3$ per group, where each case is an average of 3 tissue sections); *p*-values by *t*-test 2-tailed, * $p < 0.001$ (MBP in *C*) and * $p < 0.01$ (CC1 in *D*).

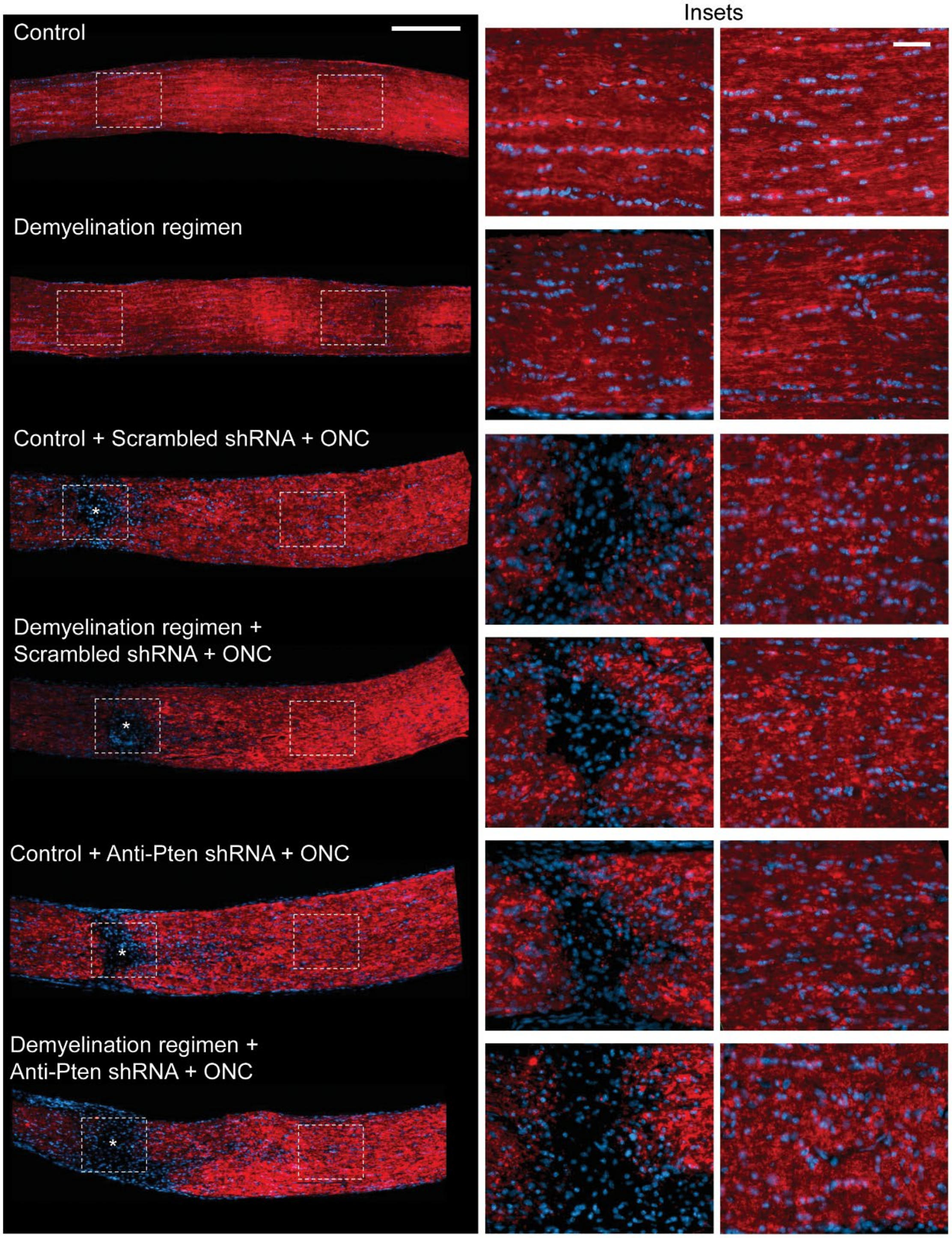


**Supplementary Figure 3. Myelin in uninjured and injured optic nerves after demyelination regimen.** Representative images of longitudinal optic nerve sections immunostained for MBP and DAPI 2 weeks after ONC or uninjured, with or without demyelination regimen, across various conditions as marked (experimental timeline in Fig. 1A). ★ - indicates crush site. Insets: Images of the uninjured or injured site, and of distal regions, are magnified for better visualization of MBP signal and DAPI-labeled nuclei. Scale bars, 200 μm (main panels), 50 μm (insets).

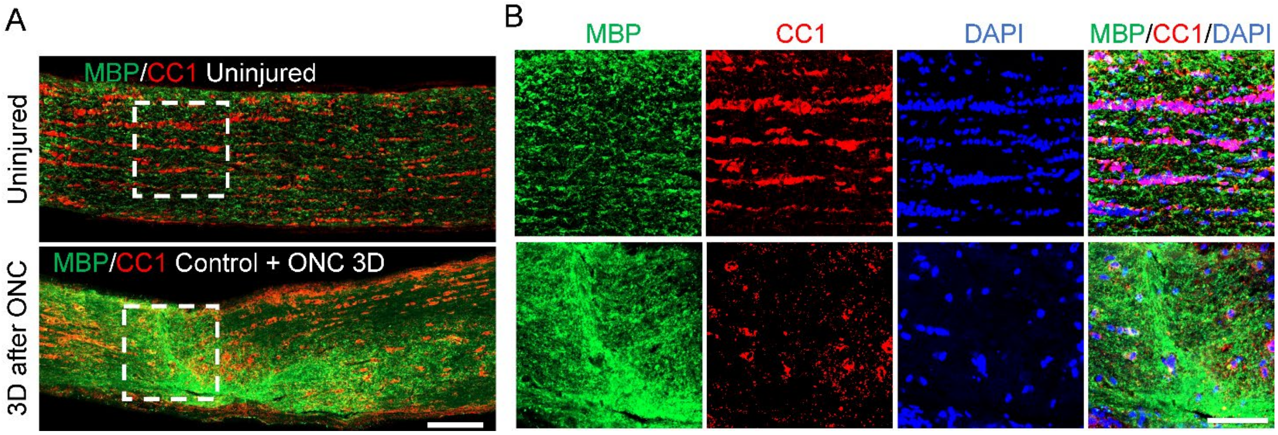


**Supplementary Figure 4. Loss of oligodendrocytes and accumulation of myelin debris in the optic nerve injury site at 3 days after injury.** (**A-B**) Representative images of uninjured (upper panels) and injured (3 days after ONC; lower panels) optic nerves longitudinal sections immunostained for CC1 (mature oligodendrocyte marker), MBP (myelin marker), and DAPI (nuclear marker), show substantial decrease in overall cellular population (fewer DAPI+ cells), including the loss of oligodendrocytes (fewer CC1+ cells), and an accumulation of myelin debris from dead oligodendrocytes (MBP+ signal) in the injury site at 3 days after ONC (lower panels in *A*-*B*), compared to uninured equivalent region of the optic nerve (upper panels in *A*-*B*). Injury site, or an equivalent region in the uninjured optic nerve, outlined with dashed white lines box in *A* and shown in insets in *B*. Scale bars: 100 µm main panels (*A*), 50 µm insets (*B*).

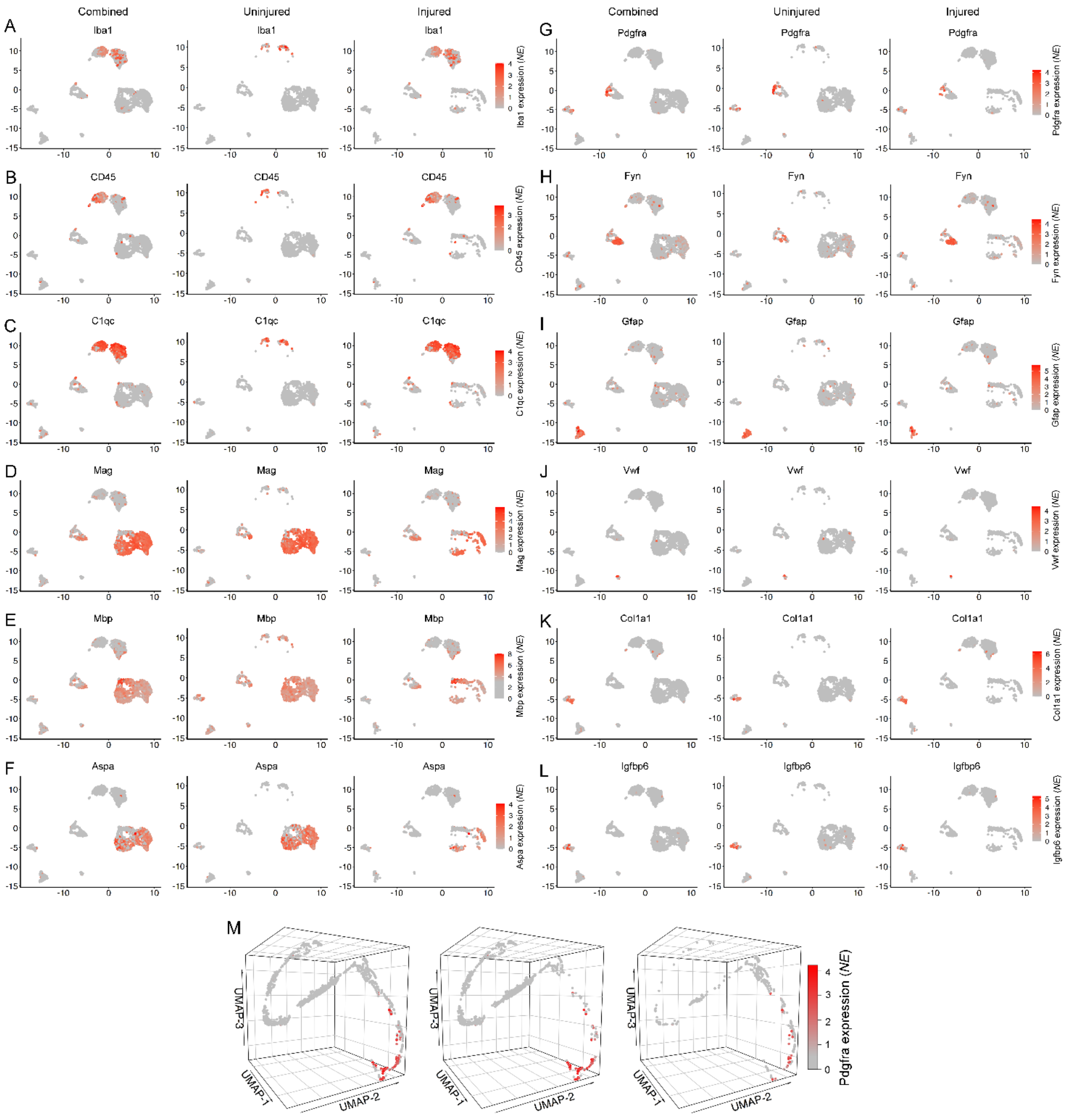


**Supplementary Figure 5. Cell type markers in single cell transcriptome profiling of uninjured and injured optic nerves.** (**A-L**) 2D UMAPs of the scRNA-seq-profiled cells comprising the injured and uninjured optic nerve tissues (with all with all cells combined, or uninjured only, or injured only), showing the expression of respective cell types' gene markers (as marked): Immune cell (Ida1, CD45, and C1qc, in *A-C*), oligodendrocyte (MAG, MBP, and Aspa, in *D-F*), oligodendrocyte progenitor cell (OPC; Pdgfra, in *G*), newly formed oligodendrocyte (NFO; Fyn, in *H*), astrocyte (GFAP, in *I*), endothelial cell (Vwf[89], in *J*), and pericyte/fibroblast (Col1a1[54] and Igfbp6[54], in *K-L*). Scale bar, color-coded normalized expression (*NE*). (**M**) 3D UMAPs of cells comprising oligodendrocyte lineage (as in Fig. 2 *F-L*; with all cells combined, or uninjured only, or injured only), showing expression of the OPC gene marker, Pdgfra (due to space limit in Fig. 2, shown in the supplementary information). Scale bar, color-coded normalized expression (*NE*).

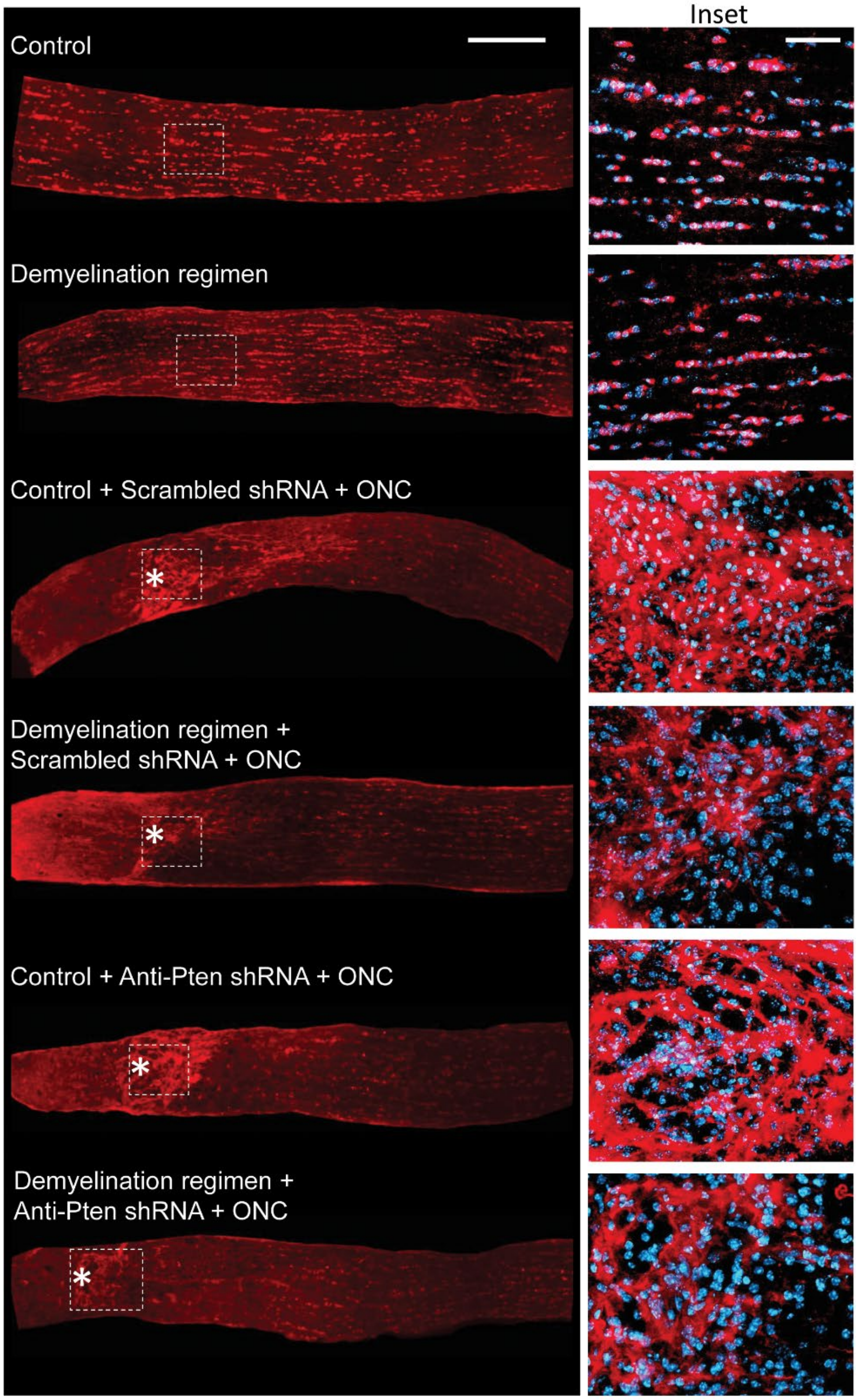


**Supplementary Figure 6. Mature oligodendrocytes' soma in uninjured and injured optic nerves after demyelination regimen.** Representative images of the longitudinal optic nerve sections immunostained for CC1 and DAPI 2 weeks after ONC or uninjured, with or without demyelination regimen, across various conditions as marked (experimental timeline in Fig. 1A). ⁎ - indicates crush site. Insets: Images of the uninjured or injured sites are magnified for better visualization of CC1 signal and DAPI-labeled nuclei. Scale bars, 200 μm (main panels), 50 μm (insets).

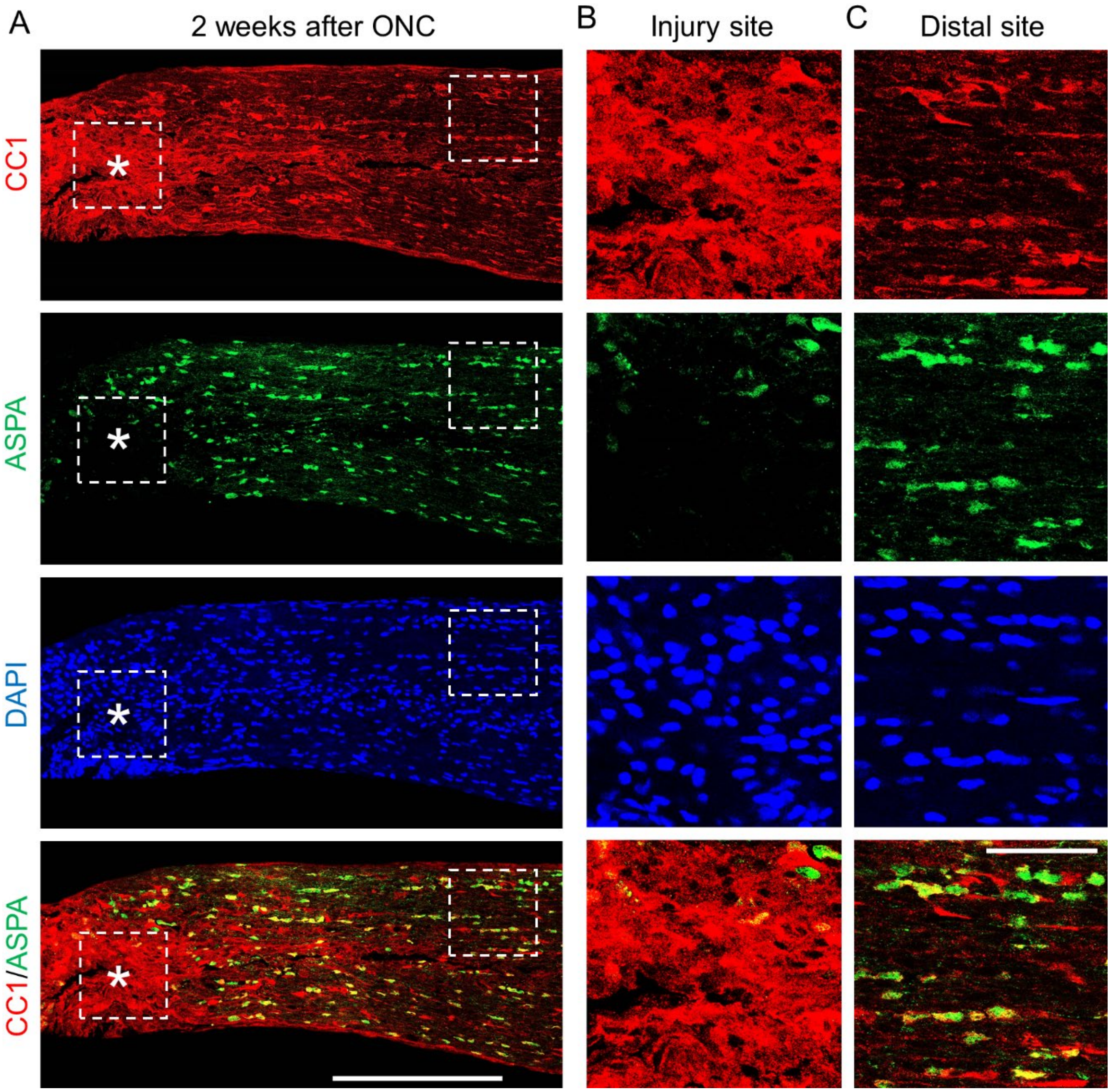


**Supplementary Figure 7. CC1+ oligodendrocytes in the injury site do not express Aspa**. (**A**) Representative confocal images of the longitudinal optic nerve section immunostained for CC1 and Aspa oligodendrocyte markers 2 weeks after ONC. ⋆ - indicates crush site. (**B**-**C**) Insets (outlined with dashed lines in *A*) magnified for better visualization of Aspa, CC1, and DAPI in the injured crush site (*B*, inset to the left in *A*), and uninjured distal from crush site region (*C*, inset to the right in *A*). Compared to the uninjured region, in the crush site CC1 density is increased whereas Aspa density is decreased, with CC1+/Aspa+ cells predominantly in the uninjured region and CC1+/Aspa- cells predominantly in the injury site. Scale bars, 200 μm (main panel), 50 μm (insets).

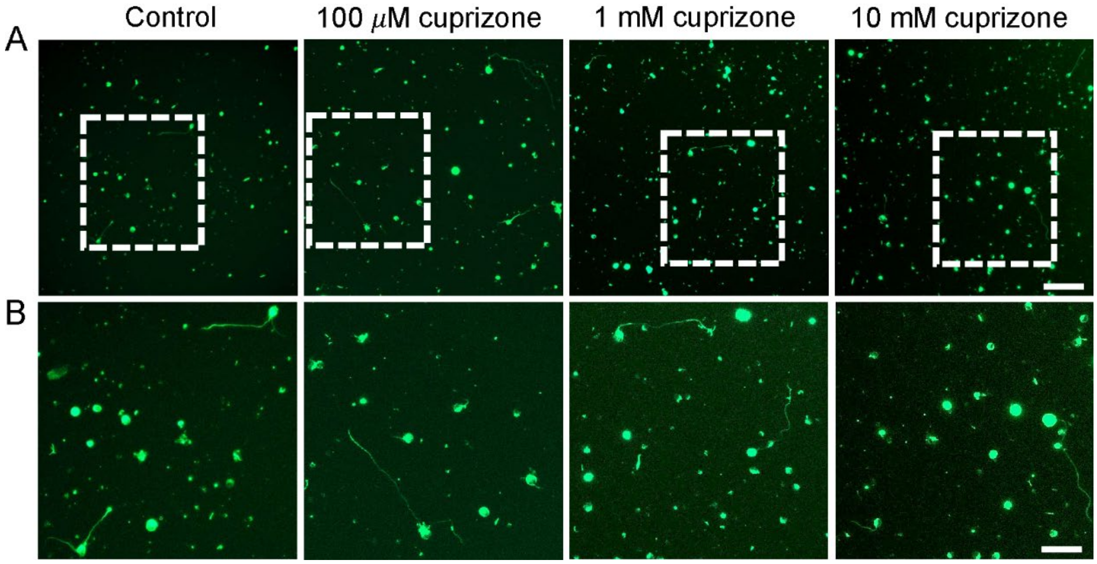


**Supplementary Figure 8. RGCs incubated with varying concentrations of cuprizone.** Adult RGCs, isolated by immunopanning for Thy1, immunostained with neuronal marker βIII-Tubulin after 5 days in culture in a defined growth medium (see Methods) with varying concentrations of cuprizone, as indicated. (**A**-**B**) There was no apparent difference in RGC density (larger field of view in *A*) or axon growth (insets in *B*) between control or cuprizone treated conditions. Scale bars, 100 μm (main panels), 50 μm (insets).
